# Tumor suppressor let-7 acts as a key regulator for maintaining pluripotency gene expression in Muse cells

**DOI:** 10.1101/2023.08.24.554727

**Authors:** Gen Li, Shohei Wakao, Masaaki Kitada, Mari Dezawa

## Abstract

In embryonic stem cells (ESCs) and induced pluripotent stem cells (iPSCs), the expression of an RNA-binding pluripotency-relevant protein, LIN28, and the absence of its antagonist, the tumor-suppressor microRNA (miRNA) let-7, play a key role in maintaining pluripotency. Muse cells are non-tumorigenic pluripotent-like stem cells residing in the bone marrow, peripheral blood, and organ connective tissues as pluripotent surface marker SSEA-3(+). They express pluripotency genes, differentiate into triploblastic-lineage cells, and self-renew at the single cell level. Muse cells do not express LIN28 but do express let-7 at higher levels than in iPSCs. In Muse cells, we demonstrated that let-7 inhibited the PI3K-AKT pathway, leading to sustainable expression of the key pluripotency regulator KLF4 as well as its downstream genes, *POU5F1*, *SOX2*, and *NANOG*. Let-7 also suppressed proliferation and glycolysis by inhibiting the PI3K-AKT pathway, suggesting its involvement in non-tumorigenicity. Furthermore, the MEK/ERK pathway is not controlled by let-7 and may have a pivotal role in maintaining self-renewal and suppression of senescence. The system found in Muse cells, in which the tumor suppressor let-7, but not LIN28, tunes the expression of pluripotency genes, might be a rational cell system conferring both pluripotency-like properties and a low risk for tumorigenicity.

## Introduction

The RNA-binding protein LIN28 and micro RNA (miRNA) lethal-7 (let-7) family members were first discovered in *Caenorhabditis elegans* (*C. elegans*) and are conserved across species (1–5). In mammals, LIN28 and let-7 work as a mutually antagonistic system. LIN28 binds to the loop of precursor-let-7 (pre-let-7) to inhibit its maturation, and conversely, let-7 binds to the 3’ untranslated region (UTR) of *LIN28* to inhibit its translation (6, 7). In this manner, the LIN28-let-7 axis widely controls many biologic processes, such as stem cell maintenance, development, differentiation, and cellular metabolism (8–11). For example, LIN28 is highly expressed during early embryogenesis but its expression declines during development (12). In mouse zygotes, LIN28 knockdown induces arrest between the 2- and 4-cell-stages, leading to a developmental failure at the morula and blastocyst stages, which suggests its importance in early development (13). LIN28 and pluripotent factors POU5F1, SOX2, and NANOG are sufficient to convert human fibroblasts into induced pluripotent stem cells (iPSCs) (14), suggesting that LIN28 is a key regulator for controlling pluripotency. The LIN28-let-7 axis works in a seesaw manner to maintain the balance between self-renewal and differentiation in pluripotent stem cells (PSCs) such as embryonic stem cells (ESCs) and iPSCs; the expression of LIN28 is high in undifferentiated ESCs and iPSCs and decreases during cell differentiation (4, 15, 16), while expression of the let-7 family, which is not observed in undifferentiated ESCs and iPSCs, increases during differentiation.

After birth, most somatic cells lose the expression of LIN28. The recurrence of LIN28 expression, however, is observed in many human cancers, such as breast-, colon-, liver-, and ovarian cancers (17). LIN28 is considered a marker for cancer stem cells and could be a target for anticancer therapies (18–20). Therefore, LIN28 is considered an oncogene. In contrast, let-7 is downregulated in many cancer cells (21–23), and forced expression of let-7 induces a slowdown of tumor growth (24). Therefore, let-7 is considered a tumor suppressor miRNA. Thus, the high expression of let-7 in the majority of somatic cells might be a strategy to decrease tumorigenic risk.

Multilineage differentiating stress-enduring (Muse) cells are endogenous reparative pluripotent-like stem cells that reside in the bone marrow, peripheral blood, and organ connective tissue as cells positive for a pluripotency surface marker, stage-specific embryonic antigen (SSEA)-3 (25–28). Muse cells are also collectible as several percent of SSEA-3(+) cells from cultured mesenchymal stromal cells (MSCs) and fibroblasts. Muse cells express other pluripotency markers, including NANOG, POU5F1, and SOX2, at moderate levels compared with ESCs and iPSCs (26, 29). They can generate endodermal-, mesodermal-, and ectodermal-lineage cells and self-renew at the single cell level; they also exhibit stress tolerance due to a high capacity for sensing and repairing DNA damage (26, 30, 31). Unlike ESCs and iPSCs, however, Muse cells are non-tumorigenic, consistent with the fact that they are endogenous to the body; express telomerase at a low level, comparable to that in somatic cells such as fibroblasts; and do not form teratomas after transplantation in vivo (26). Circulating endogenous Muse cells and intravenously administered exogenous Muse cells both selectively home to damaged tissue by sensing sphingosine-1-phosphate, a damage signal produced by damaged tissue; phagocytose apoptotic differentiated cell fragments to receive differentiation machineries such as transcription factors; initiate differentiation into the same cell type as the phagocytosed cells in a short time period; and repair the tissue by replacing damaged cells (28, 32–36). As demonstrated in a rabbit acute myocardial infarction model, allogeneic Muse cells can escape host immunologic attack and survive as functional cells in the host tissue for more than half a year without immunosuppression. The immune privilege of Muse cells is partly explained by the expression of human leukocyte antigen (HLA)-G, which is expressed in extravillous trophoblast cells in the placenta and plays an important role in immune tolerance during pregnancy (35). Clinical trials are currently being conducted for stroke, acute myocardial infarction, epidermolysis bullosa, spinal cord injury, amyotrophic lateral sclerosis, and COVID19-acute respiratory distress syndrome using intravenous injections of human clinical-grade Muse cells without HLA-matching or immunosuppression. The safety and effectiveness of Muse cells have been reported in clinical trials of acute myocardial infarction and epidermolysis bullosa (37, 38).

To clarify the mechanisms underlying how Muse cells maintain their pluripotency-like properties without being tumorigenic, we evaluated the LIN28-let-7 axis in Muse cells. We found that Muse cells did not express LIN28 but expressed let-7 at higher levels than iPSCs. We also explored the mechanisms by which let-7 acts as a key regulator of pluripotency gene expression and its involvement in suppressing proliferation and glycolysis in Muse cells.

## Results

### Expression of LIN28 and let-7 in Muse cells

We isolated SSEA-3(+)-Muse cells from human bone marrow (BM)-MSCs and normal human dermal fibroblasts (NHDFs) as reported previously (Supplemental Fig. S1A and S1B) (26, 39). When the BM-MSCs were cultured with fibroblast growth factor 2 (FGF2) for 2 population doubling levels (PDLs), the ratio of the Muse cell population among MSCs was 4.82% on average from 3 replicates, similar to previous reports (39) (Figure 1A). When the BM-MSCs were cultured without FGF2, the Muse cell ratio decreased to 1.22% on average from 3 replicates (Figure 1A). Therefore, FGF2 is indispensable for maintaining Muse cells in BM-MSC populations.

**Figure 1.**
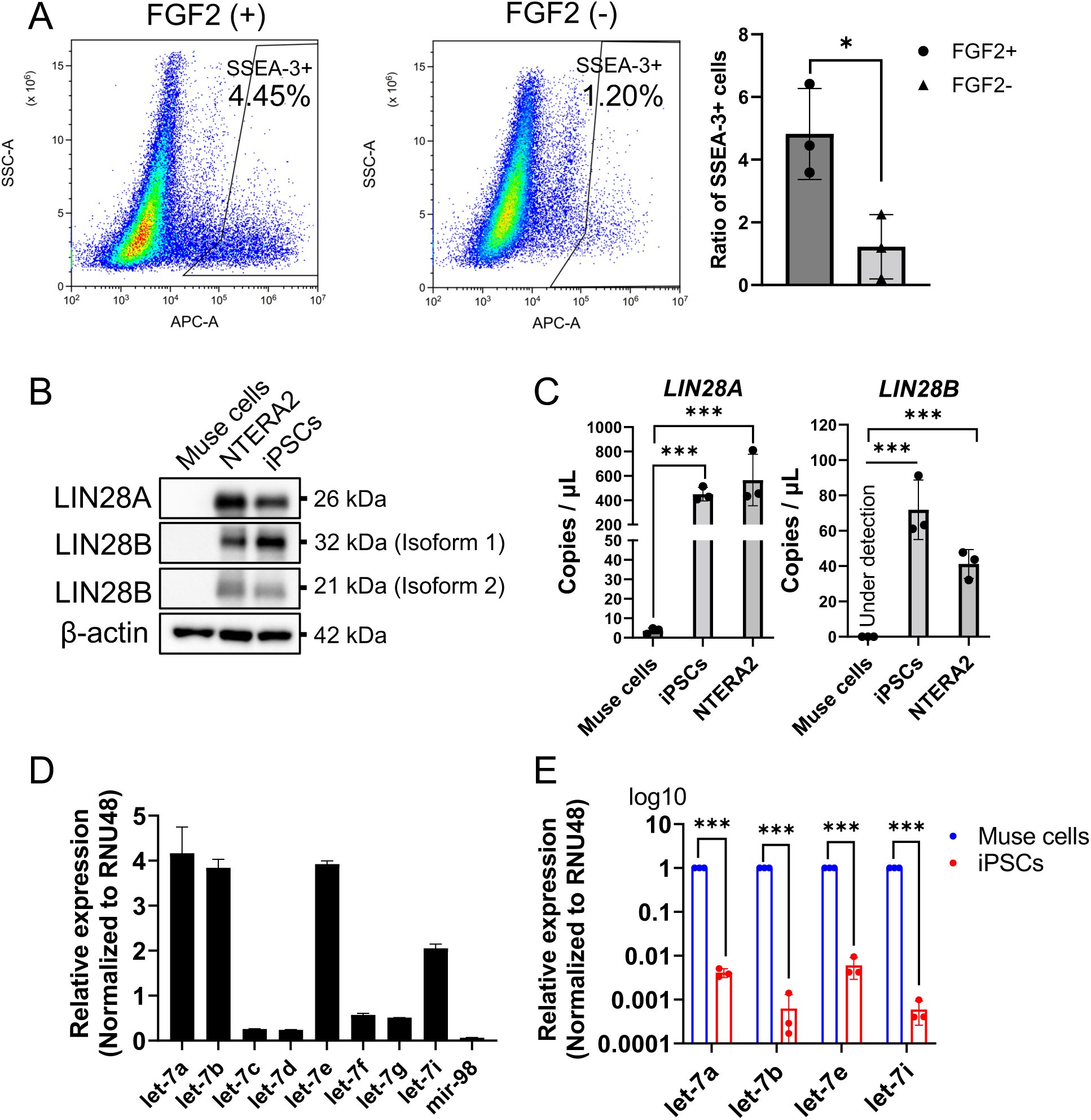
Muse cells express let-7 but not LIN28. (A) The ratio of the Muse cell population, which was cultured with or without FGF2 treatment (n=3). (B) Western blot of LIN28A/B in Muse cells, NTERA2, and iPSCs. β-Actin was used as an endogenous control. (C) Expression of *LIN28A* and *LIN28B* in Muse cells, NTERA2, and iPSCs in ddPCR. Data are shown as DNA copies/μL (all, n=3). (D) Expression of let-7 subtypes in BM-Muse cells in qPCR (all n=3). RNU48 was used as an endogenous control. (E) qPCR of let-7a, -7b, -7e, and -7i in Muse cells and iPSCs (n=3). RNU48 was used as an endogenous control. A log10 scale was used for the y-axis.

In Western blotting, both LIN28A and LIN28B were under the detection limit in Muse cells, while they were detectable in the human teratoma cell line NTERA2 and human iPSCs (Fig. 1B). In droplet digital PCR (ddPCR), the expression of *LIN28A* in Muse cells was significantly lower than that in iPSCs and NTERA2; the expression of *LIN28B* was under the detection limit in Muse cells, but clearly detectable in iPSCs and NTERA2 (Fig. 1C). Quantitative polymerase chain reaction (qPCR) showed that the major let-7 subtypes expressed in Muse cells were let-7a, -7b, -7e, and -7i among the 9 subtypes (Fig. 1D). The expression of let-7a, -7b, -7e, and -7i was 100–1000 times higher in Muse cells than in iPSCs (Fig. 1E). These findings indicated that, in contrast to iPSCs, Muse cells basically express let-7 but not LIN28A/B. Muse cells isolated from NHDFs showed similar trends of LIN28A/B and let-7 expression (Supplemental Fig. S1C, S1D, S1E, and S1F). We used BM-MSC–derived Muse cells in the following experiments and focused on the let-7a, let-7b, let-7e, and let-7i subtypes.

### Gene expression profile in Muse cells after let-7 knockdown

A sequence-specific miRNA inhibition system, tough decoy (TuD) (40), was constructed and introduced into Muse cells by lentivirus for the loss of function of let-7 (Figs. 2A and S2A). We performed a luciferase assay to confirm the activity of TuD-based let-7 knockdown (KD). We first constructed 4 luciferase-expression plasmids with the target sequences of let-7a, -7b, -7e, and -7i inserted into the 3’UTR of firefly luciferase: pFluc-let-7a, pFluc-let-7b, pFluc-let-7e, and pFluc-let-7i, respectively. We also prepared 6 types of Muse cells; non-transfected naïve Muse cells, negative control-TuD-ath-mir416 introduced Muse cells (control-TuD-Muse cells); and let-7a-KD, let-7b-KD, let-7e-KD, and let-7i-KD Muse cells. The plasmids were each introduced into the 6 types of Muse cells (Fig. S2B). We measured the luciferase intensity. When the pFluc-let-7a plasmid was transfected into each type of Muse cell, the luminescence signal of the luciferase was significantly increased in the let-7a-KD-, let-7b-KD, let-7e-KD, and let-7i-KD-Muse cells compared with naïve and control-TuD-Muse cells. A similar tendency was confirmed for pFluc-let-7b, -7e, and -7i (Fig. 2B). Thus, the TuD-let-7 system effectively knocked down let-7a, -7b, -7e, and -7i expression in Muse cells, although the inhibition specificity was not high among these 4 subtypes, probably due to sequence similarities among the subtypes (Fig. S2C). Let-7a-KD-Muse cells were collected from BM-MSCs after 3∼4 PDLs after introducing the TuD-let-7a lentivirus. TUNEL staining results showed that the apoptotic cell ratio was not largely increased by transfecting Muse cells with the TuD-let-7a lentivirus (Fig. S2D). Similar trends were confirmed with the TuD-let-7b, -let-7e, and -let-7i lentivirus transfected cells (data not shown).

**Figure 2.**
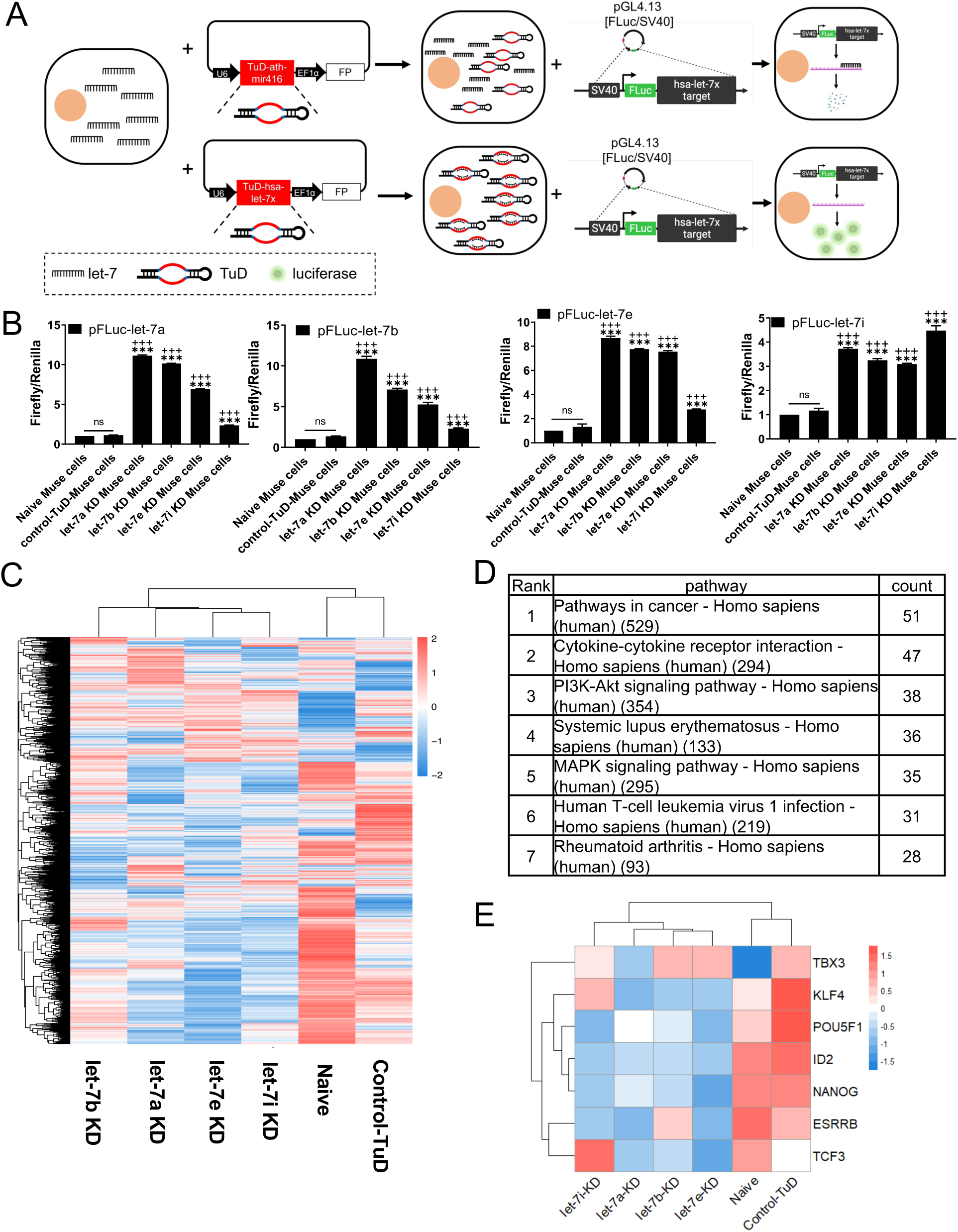
let-7 TuD knockdown and bioinformatic analysis by microarray. (A) Experimental design of TuD-based let-7 KD and evaluation of let-7 KD by luciferase assay. FP (Fluorescent protein): either EGFP or mCherry, used as an indicator for transfected cells. FLuc: firefly luciferase. Let-7x represents let-7a, -7b, -7e, or -7i. (B) Luciferase assay for let-7 KD. pFLuc-let-7a, -7b, -7e, and -7i: pGL4.13 with firefly luciferase and let-7a, -7b, -7e, or -7i target sequence inserted in the 3’UTR of firefly luciferase. ***p<0.001 vs naïve Muse cells; +++p<0.001 vs control-TuD-Muse cells; ns: no significant difference. (C) Microarray heatmap of let-7a-KD, -7b-KD, -7e-KD, -7i-KD, naïve and control-TuD Muse cells. (D) KEGG pathway analysis showing the top 7 changed pathways after let-7 KD. The number in the pathway column indicates the total factor number in the pathway, and the count indicates the number of changed factors after let-7 KD. Control-TuD-Muse cells were set as the control group. Let-7a-KD, let-7b-KD, let-7e-KD, and let-7i-KD Muse cells were combined as the experimental group. (E) Heatmap showing some of the changed pluripotency-relevant genes after let-7 KD. Red: upregulated. Blue: downregulated. White: no change.

A DNA microarray analysis was conducted to assess the transcriptome changes after let-7-KD in Muse cells. Figure 2C shows a cluster heatmap illustrating that the gene expression pattern was changed in let-7a-KD-, 7b-KD-, 7e-KD-, and 7i-KD-Muse cells compared with naïve- and control-TuD-Muse cells. Because the TuD inhibition system did not specifically downregulate each of the 4 let-7 subtypes in Muse cells, we combined the microarray gene profiles of let-7a-, let-7b-, let-7e-, and let-7i-KD Muse cells into a single group and performed the Kyoto Encyclopedia of Genes and Genomes (KEGG, https://www.genome.jp/kegg/) pathway frequency analysis for comparison with the control-TuD-Muse cell group. The KEGG pathway frequency analysis revealed the top 7 changed pathways between the control-TuD- and let-7 (-7a, -7b, -7e, and -7i) groups, which included cancer, cytokine-cytokine receptor interaction, and the PI3K-AKT and MEK/ERK signaling pathways (Fig. 2D). Interestingly, let-7-KD (-7a, -7b, -7e, and -7i)-Muse cells exhibited the downregulation of pluripotency-related genes such as *KLF4*, *POU5F1*, *ID2*, *NANOG*, and *ESRRB* (Fig. 2E), and the upregulation of cell cycle-relevant genes such as Cyclin D2 (CCND2), cell division cycle 25A (CDC25A), origin recognition complex, minichromosome maintenance proteins, and retinoblastoma protein in comparison with naïve- and control-TuD-Muse cells (Supplemental Fig. S2E). In addition, *P53* was downregulated by let-7 KD (Supplemental Fig. S2E).

As the expression of let-7a was the highest among the let-7 subtypes in BM-Muse cells (Fig. 1D) and the knockdown effect of the TuD structure was not specific (Fig. 2B), the TuD-let-7a KD system was applied in the following experiments. To predict let-7a targets, we utilized the Encyclopedia of RNA Interactomes (ENCORI, http://starbase.sysu.edu.cn/), a platform that provides information on miRNA targets collated from 7 databases (i.e., PITA, RNA22, miRmap, microT, miRnada, PicTar, and TargetScan). Genes predicted as let-7a target genes by more than 4 databases were screened, and those screened genes were then further analyzed using protein-protein interactions analysis with String (https://string-db.org/) (41).

We focused on protein-protein interactions between the predicted let-7a targets and PI3K (PIK3CA, PIK3CB, PIK3CD, and PIK3CG) or MEKK/MEK/ERK (MAPK1, MAPK3, MAP2K1, MAP2K2, and MAP3K1) because these 2 pathways were listed in the top 7 pathways that were different between the control-TuD- and let-7 groups (Fig. 2D). Among the predicted let-7a target genes, insulin receptor substrate 2 (IRS2), insulin like growth factor 1 receptor (IGF1R), insulin receptor (INSR), and neuroblastoma RAS viral oncogene homolog (NRAS) were predicted to interact with both the MEK/ERK and PI3K-AKT signaling pathways (Fig. 3A and 3B). We conducted Western blot analyses to examine the expression of the pro-form of IGF1R (pro-IGF1R), the IGF1R beta subunit (IGF1Rβ), the pro-form of INSR (pro-INSR), the INSRβ subunit (INSRβ), IRS2, and NRAS. Pro-IGF1R, IGF1Rβ, and IRS2 were significantly increased in let-7a-KD-Muse cells compared with naïve- and control-TuD-Muse cells (Fig. 3C), suggesting that IGF1R and IRS2 are targets of let-7a in Muse cells. NRAS, pro-INSR, and INSRβ expression, however, did not largely change among the naïve-, control-TuD-, and let-7a-KD-Muse cells (Fig. 3C). The molecular sizes of the Western blot signals for IRS2 differed between Muse cells and the positive control-HEK293T (Fig. 3C). This difference was considered to be the phosphorylation of IRS2 in naïve-, control-TuD-, and let-7a-KD-Muse cells in the presence of serum. We therefore used a serum-free medium to culture the Muse cells to eliminate the influence of phosphorylation on the molecular weight of IRS2 and found that the size of IRS2 in Muse cells was the same as that observed in HEK293T (Fig 3D).

**Figure 3.**
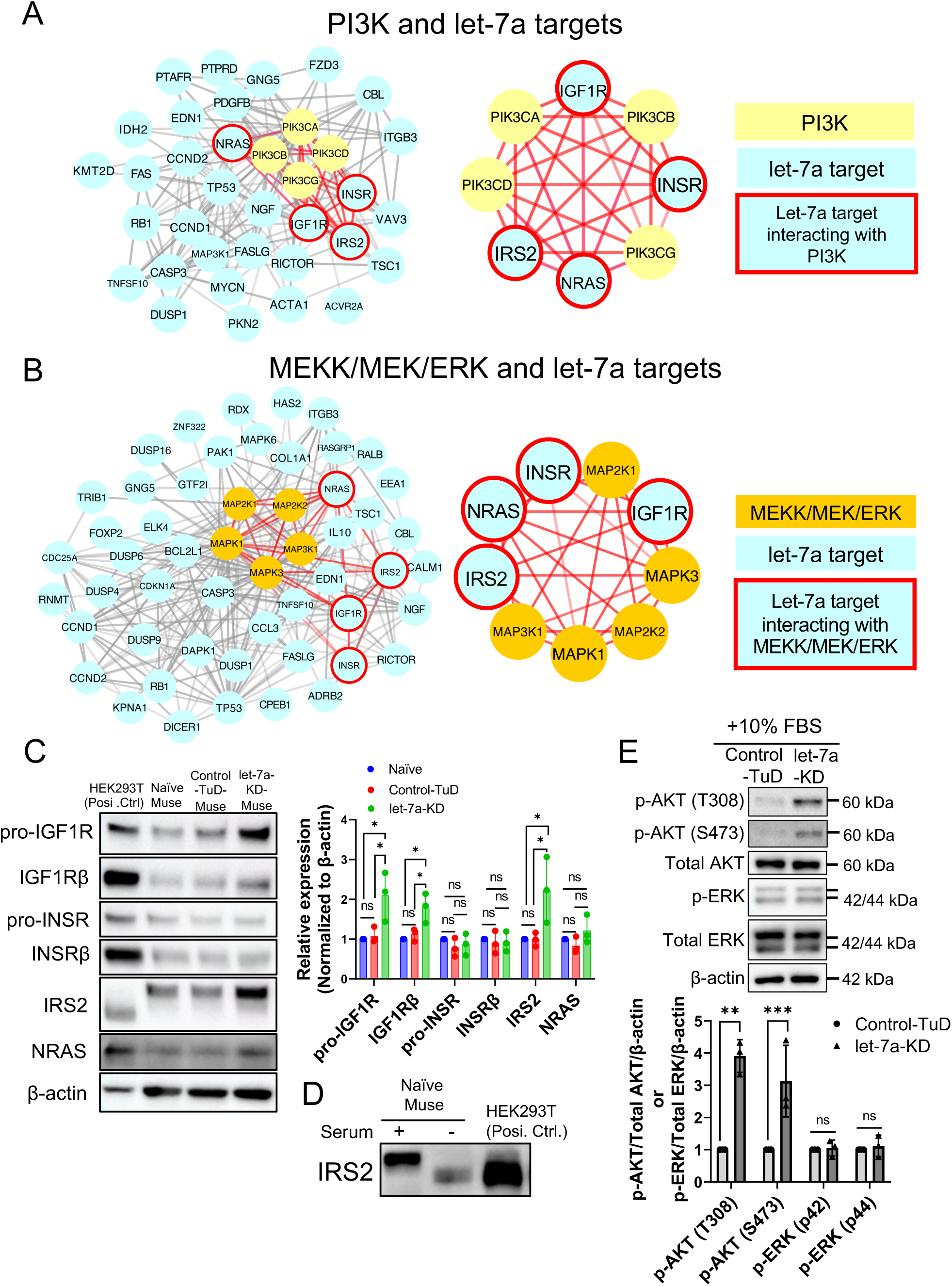
Prediction and confirmation of let-7 targets in Muse cells. (A) Protein-protein interactions of the putative let-7a targets and PI3K. Blue: let-7a targets predicted by ENCORI; Yellow: PI3K isoforms; Blue with red frame: putative let-7a targets interacting with PI3K (B) Protein-protein interactions of the putative let-7a targets and MEKK/MEK/ERK. Blue: let-7a targets predicted by ENCORI; Orange: isoforms of MEKK, MEK, or ERK; Blue with red frame: putative let-7a targets interacting with MEKK, MEK, or ERK (C) Western blot of predicted let-7a targets in Muse cells (n=3). Posi. Ctrl. (Positive control): HEK293T. Signals of each predicted target were normalized by β-actin. (D) Western blot of IRS2 under serum or serum-free conditions. Posi. Ctrl. (Positive control): HEK293T. (E) Western blot of p-AKT and p-ERK in control-TuD- and let-7a-KD-Muse cells under 10% FBS culture condition (n=3). β-Actin was used as endogenous control. p-AKT/total AKT/β-actin: The p-AKT intensity was divided by that of total AKT, and then further divided by that of β-actin to determine the quantitative value. Similarly, the p-ERK intensity was divided by that of total ERK, and then further divided by that of β-actin: P42 and p44 were calculated separately.

Activation of PI3K and its downstream cascade converge on phosphorylated AKT (p-AKT), indicating that PI3K is downstream of IGF1R/IRS2 and AKT is the critical molecule in the PI3K pathway (42). In addition, IGF1R and IRS2 are reported to be let-7 targets (43, 44). Thus, we compared the expression of p-AKT (T308), p-AKT (S473), and p-ERK (p42/p44) between control-TuD and let-7a-KD-Muse cells. In Muse cells, let-7a KD upregulated the expression of both p-AKT (T308) and p-AKT (S473), but not p-ERK (p42/p44) (Fig. 3E). Thus, let-7 inhibited the PI3K-AKT pathway by inhibiting IGF1R and IRS2, but had a limited influence on the MEK/ERK pathway.

### Effect of let-7 on PI3K-AKT and crosstalk between the MEK/ERK and PI3K-AKT pathways

We examined how let-7-KD affected the PI3K-AKT pathway in Muse cells. To confirm activation of the PI3K pathway, we cultured naïve-, control-TuD-, and let-7a-KD-Muse cells for 8 h under serum-free conditions to remove the influence of phosphorylation by the growth factors contained in the serum. In the serum-free conditions, p-AKT (T308) and p-AKT (S473) were not detected in any of the 3 types of Muse cells (i.e., naïve, control-TuD, and let-7a-KD) on Western blots (Fig. 4A). After adding 100 ng/mL insulin to the serum-free medium for 15 min, the p-AKT (T308) and p-AKT (S473) levels increased in all 3 types of Muse cells. The highest increase of p-AKT (T308) and p-AKT (S473) was in let-7-KD-Muse cells compared with naïve- and control-TuD-Muse cells (Fig. 4B). A similar trend was observed after the addition of 100 ng/mL IGF1 to the serum-free culture medium for 15 min. Notably, the increased p-AKT (T308) and p-AKT (S473) levels were higher in 100 ng/mL IGF1-treated naïve, control-TuD, and let-7a-KD Muse cells than in 100 ng/mL insulin-treated cells (Fig. 4B and 4C).

**Figure 4.**
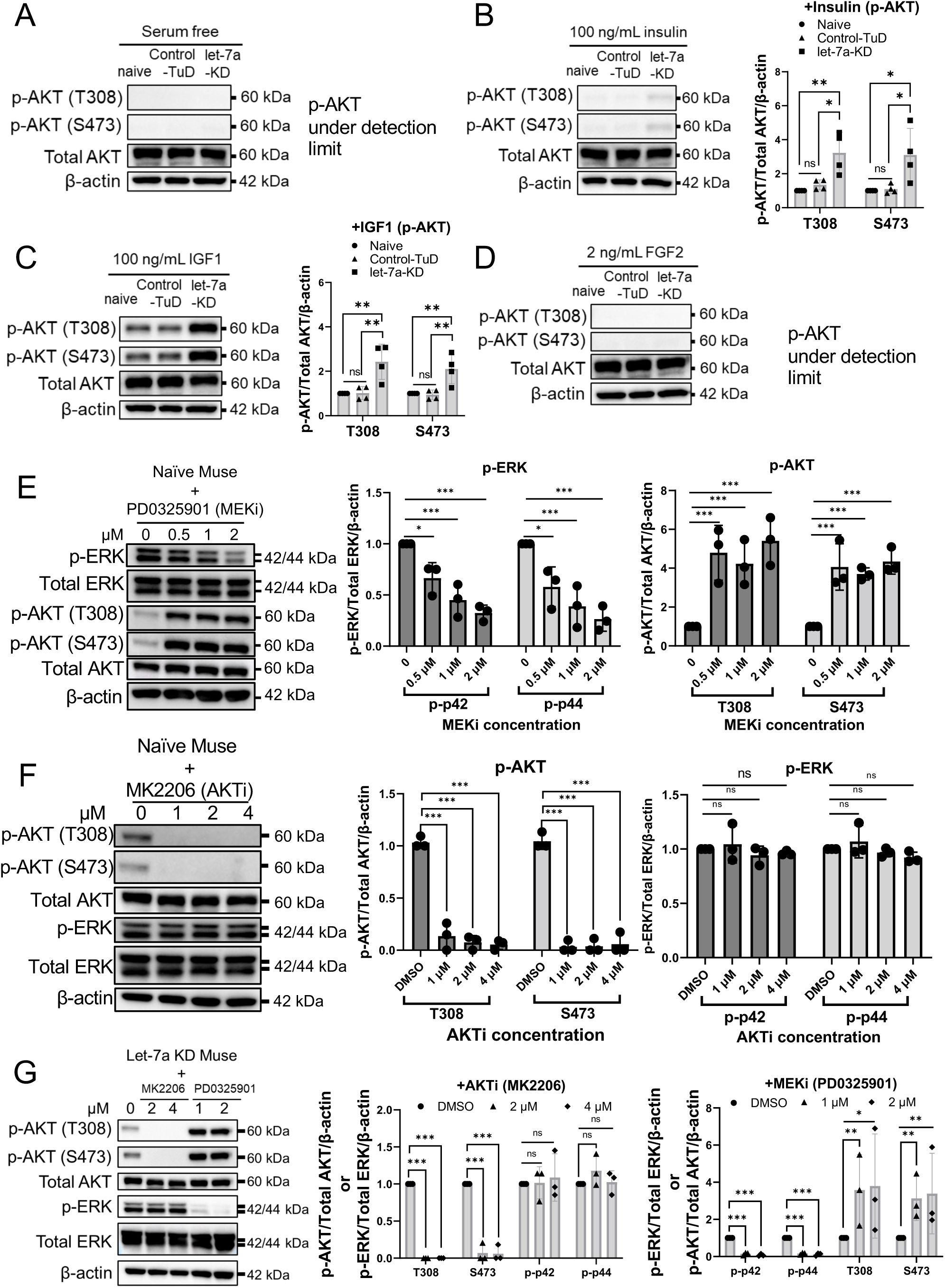
Effect of let-7 on PI3K-AKT and crosstalk between the MEK/ERK and PI3K-AKT. (A-D) Serum starvation eliminated AKT phosphorylated sites in naïve, control-TuD-, and let-7-KD-Muse cells. Adding insulin (100 ng/mL) or IGF1 (100 ng/mL) phosphorylated AKT at T308 and S473 sites, and p-AKT increased more in let-7-KD Muse cells than in naïve and control-TuD-Muse cells (all n=4). Addition of 2 ng/mL FGF2 did not change the p-AKT level. (E) In naïve Muse cells, MEKi (PD0325901) inhibited p-ERK (p-p42/p-p44) in a dose-dependent manner. Inhibition of p-ERK increased the level of p-AKT (T308) and p-AKT (S473) (all, n=3). (F) In naïve Muse cells, AKTi (MK2206) inhibited p-AKT (T308) and p-AKT (S473), but the expression of p-ERK (p-p42/p-p44) was not affected (all, n=3). (G) In let-7a-KD Muse cells, the addition of AKTi (MK2206) and MEKi (PD0325901) inhibited p-AKT and p-ERK, respectively. MEKi increased p-AKT, but AKTi did not upregulate p-ERK. β-Actin was used as endogenous control. p-AKT/total AKT/β-actin: the p-AKT intensity was divided by that of total AKT, and then further divided by that of β-actin to determine the quantitative value. Similarly, the p-ERK intensity was divided by that of total ERK, and then further divided by that of β-actin: P42 and p44 were calculated separately.

FGF2 is important for maintaining Muse cells, as suggested in Fig. 1A. Therefore, the effect of FGF2 on the PI3K-AKT pathway was examined. The increased AKT phosphorylation, however, remained under the detection limit when 2 ng/mL FGF2 was supplied to the serum-free culture medium for 15 min (Fig. 4D). Thus, let-7a-KD activated the PI3K-AKT pathway in the presence of insulin or IGF1, but not FGF2, in Muse cells.

The interaction between the MEK/ERK and PI3K-AKT pathways was investigated in naïve Muse cells. PD0325901, a MEK inhibitor (MEKi), induced a dose-dependent decrease in p-ERK. The p-AKT (T308) and p-AKT (S473) levels, however, were both upregulated in naïve Muse cells (Fig. 4E). Treating Muse cells with MK2206, an AKT inhibitor (AKTi), inhibited the phosphorylation of AKT at T308 and S473, but did not affect p-ERK (Fig. 4F). These results suggested that, in naïve Muse cells, MEK/ERK inhibited the PI3K-AKT pathway by inhibiting the phosphorylation of AKT, while AKT inhibition did not affect the phosphorylation of ERK.

In let-7a-KD-Muse cells, MEKi also upregulated p-AKT (T308) and p-AKT (S473), while AKTi-induced inhibition of p-AKT did not affect p-ERK expression, similar to the findings in naïve Muse cells (Fig. 4G). These results suggested that, although let-7a KD changed the transcriptome of Muse cells, the crosstalk between the PI3K-AKT and MEK/ERK pathways was not largely affected.

### Let-7 KD decreased the expression of pluripotency genes

We treated BM-MSCs with LY294002, a PI3K inhibitor (PI3Ki), and MEKi for 2 PDLs, and examined the Muse cell population ratio. The results of 3 replicates showed that, compared with the DMSO-treated group, PI3Ki treatment did not largely affect the Muse cell ratio, while MEKi significantly reduced the Muse cell ratio to less than 1% (Fig. 5A). These results suggested that the MEK/ERK pathway, but not the PI3K-AKT pathway, has a pivotal role in maintaining Muse cells.

**Figure 5.**
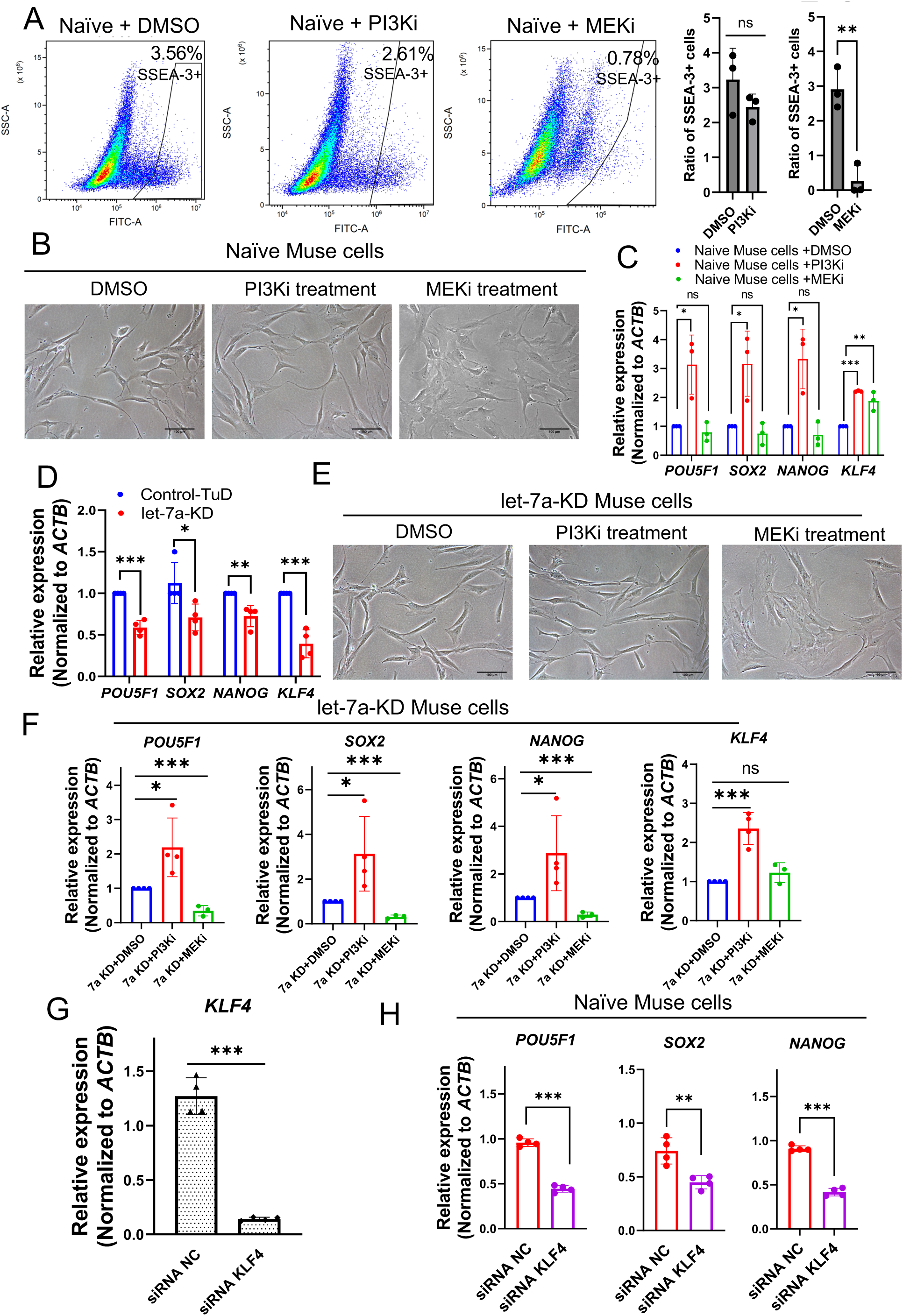
let-7 promoted the expression of pluripotency genes. (A) Flow cytometry analysis of Muse cells in DMSO-, PI3Ki-, and MEKi-treated hMSCs. MEKi decreased the ratio of Muse cells, but PI3Ki showed no significant effect (n=3). (B) Cell morphology of naïve Muse cells 24 h after DMSO, PI3Ki, and MEKi treatment. Scale bar: 100 µm (C) Comparison of pluripotency gene expression among DMSO-, PI3Ki-, and MEKi-treated naïve Muse cells by qPCR (all n=3). (D) Expression of pluripotency genes between control-TuD and let-7a-KD Muse cells in qPCR (all n=4). (E) Cell morphology of let-7a-KD Muse cells 24 h after DMSO, PI3Ki, and MEKi treatment. Scale bar: 100 µm (F) qPCR of *POU5F1*, *SOX2*, and *NANOG* in DMSO- (n=4), PI3Ki- (n=4), and MEKi-treated (n=3) let-7a-KD Muse cells. (G) qPCR of *KLF4* in negative control (NC) siRNA and KLF4 knockdown siRNA (n=4). (H) qPCR of *POU5F1*, *SOX2*, and *NANOG* in negative control (NC) siRNA and KLF4-knockdown siRNA-introduced naïve Muse cells (n=4). PI3Ki: LY294002. MEKi: PD0325901. ACTB was used as an endogenous control in all the qPCR experiments.

We then examined how *POU5F1*, *SOX2*, *NANOG*, and *KLF4* were affected by PI3Ki and MEKi treatment in naïve Muse cells. The morphology of naïve Muse cells did not remarkably change after PI3Ki treatment for 24 h, while the morphology of MEKi-treated Muse cells became flattened (Fig. 5B). PI3Ki treatment induced a significantly higher (2 to 4-fold) expression of *POU5F1*, *SOX2*, *NANOG*, and *KLF4* in naïve Muse cells compared with DMSO-treated naïve Muse cells (Fig. 5C). MEKi incubation for 24 h, however, did not change the *POU5F1*, *SOX2*, and *NANOG* expression compared with that in DMSO-treated naïve Muse cells, but increased the expression of *KLF4* by ∼2 times (Fig. 5C).

Following differentiation of ESCs, pluripotency gene expression decreases and let-7 expression increases (45, 46). In contrast to ESCs, let-7a-KD significantly decreased *POU5F1*, *SOX2*, *NANOG*, and *KLF4* expression in Muse cells compared with control-TuD-Muse cells (Fig. 5D), consistent with the microarray data (Fig. 2E). We then treated TuD-let-7a-KD Muse cells with PI3Ki or MEKi for 24 h. PI3Ki-treated let-7a-KD Muse cells did not show clear changes in cell morphology. In contrast, the morphology of MEKi-treated let-7a-KD Muse cells became flattened (Fig. 5E). The qPCR results showed that treating let-7a-KD Muse cells with PI3Ki for 24 h reversed the effects of let-7a-KD; expression of *POU5F1*, *SOX2*, *NANOG*, and *KLF4* was significantly elevated compared with that in control (DMSO supplied-let-7a-KD) Muse cells (Fig. 5F). This finding suggested that let-7 controlled the expression of these genes through the PI3K-AKT pathway.

MEKi treatment of let-7a-KD for 24 h significantly suppressed the expression of *POU5F1*, *SOX2*, and *NANOG* compared to DMSO-treated let-7a-KD Muse cells (Fig. 5F). Due to the morphology changes after MEKi treatment, we considered that the decrease in *POU5F1*, *SOX2*, and *NANOG* expression might relate to cell senescence or apoptosis. To investigate this, we cultured control-TuD- and let-7a-KD-Muse cells with MEKi for 5 days. Flow cytometry analysis of SPiDER β-gal staining showed that MEKi treatment nonsignificantly increased senescence-associated beta-galactosidase (SA-βgal) intensity ∼1.5 times more in control-TuD Muse cells than in DMSO-treated Muse cells, whereas it significantly (∼2-fold) increased SA-βgal intensity in let-7a-KD Muse cells compared with DMSO-treated Muse cells on days 1, 3, and 5 (Supplemental Fig. S3A). PI3Ki treatment did not increase the fluorescence intensity in either the control-TuD- or let-7a-KD-Muse cells (Supplemental Fig. S3B). The flow cytometry results of AnnexinV-FITC staining showed that neither PI3Ki nor MEKi induced severe apoptosis compared with the UV-treated positive control (Supplemental Fig. S3C). These findings suggested that MEKi induced cell senescence rather than apoptosis in let-7a-KD Muse cells and that let-7a inhibits cellular senescence when the MEK/ERK pathway is blocked.

KLF4 controls the expression of SOX2 and NANOG in ESCs (47–49). To examine whether KLF4 regulates the expression of *POU5F1*, *SOX2*, and *NANOG* in Muse cells, *KLF4* KD was conducted using small interfering RNA (siRNA). Western blotting showed a sharp decrease in KLF4 in Muse cells at 2 and 3 days after transfection of 3 different siRNAs (siRNA #1, siRNA #2, and siRNA #3; Supplemental Fig. S3D). When the 3 siRNAs for *KLF4* were mixed, qPCR confirmed the statistically significant suppression of *KLF4* on day 3 after siRNA transfection (Fig. 5G). Consequently, *POU5F1* (p<0.001), *SOX2* (p<0.01), and *NANOG* (p<0.001) were suppressed in KLF4-KD-Muse cells compared with siRNA-negative control-introduced Muse cells (Fig. 5H). These findings suggested that KLF4 positively regulates *POU5F1*, *SOX2*, and *NANOG* expression in Muse cells.

### Overexpression of mature let-7 was not feasible in Muse cells

We attempted to overexpress mature let-7a, let-7b, let-7e, and let-7i in Muse cells by introducing pre-let-7-lentivirus (Supplemental Fig. S4A). Mature let-7a is generated from 3 kinds of pre-let-7a (Roush et al., 2008). We first compared their expression levels in naïve Muse cells. qPCR results showed that pre-let-7a-3 was dominantly expressed among the three pre-let-7a forms, corresponding to ∼7 times and ∼2 times higher than pre-let-7a-1 and pre-let-7a-2, respectively (Supplemental Fig. S4B). Therefore, we selected pre-let-7a-3 for the let-7 overexpression experiment. The pre-let-7a-3 was confirmed to be overexpressed by more than ∼15 times compared with empty vector-transfected Muse cells (Supplemental Fig. S4C). While mature let-7a was overexpressed at an average of only 1.08 times higher than that in empty vector-transfected Muse cells after culturing for 5 PDLs, however, the increase was statistically significant (Supplemental Fig. S4C). In addition, lentiviral transfection with pre-let-7b and pre-let-7e failed to overexpress mature let-7b and let-7e in naïve Muse cells (Supplemental Fig. S4C). Both pre-let-7a-3 and mature let-7a were largely overexpressed in NTERA2 cells, demonstrating that the overexpression system worked well in NTERA2 cells but not in Muse cells (Supplemental Fig. S4D).

Notably, mature let-7i was significantly overexpressed (1.5∼2 times higher than that in empty vector-transfected Muse cells) (Supplemental Fig. S4C). Luciferase assay confirmed that let-7i overexpression inhibited the expression of firefly luciferase compared with that in pGL4.13-empty vector-transfected cells (Supplemental Fig. S4E). We then examined the influence of let-7i overexpression on pluripotency gene expression. The expression of *POU5F1*, *SOX2*, *NANOG*, and *KLF4* did not significantly change in the let-7i overexpressing Muse cells compared with empty vector-transfected cells (Supplemental Fig. S4F). One possible explanation is that the effect of the let-7 family members on positive regulation of pluripotency genes had already reached a plateau prior to their overexpression.

### let-7a KD accelerated Muse cell proliferation without activation of telomerase

Cell cycle analysis by flow cytometry revealed that the percentage of naïve-Muse cells at the G0/G1, S, and G2/M phase was 93%, 4.4%, and 2.5% at 0 h, respectively (Fig. 6A). After seeding the cells on culture dishes, naïve Muse cells started entering S phase at 24 h. Control-TuD-Muse cells showed a similar trend. In let-7a-KD-Muse cells, the G0/G1, S, and G2/M percentages were similar to those in naïve- and control-TuD-Muse cells at 0 h. The let-7a-KD-Muse cells, however, started entering into S phase after 12 h, which was earlier than that of the naïve- and control-TuD-Muse cells (Fig. 6A). The proliferation speed of the control-TuD- and let-7a-KD-Muse cells was examined for 6 days. Let-7a-KD-Muse cells had higher proliferation activity than control-TuD-Muse cells starting on day 3 (p<0.01), and the growth rate was 2.3 times higher in let-7a-KD-Muse cells than in control-TuD-Muse cells (p<0.001) on day 6 (Fig. 6B). These results demonstrated that let-7a KD accelerated the cell cycle and proliferation speed in Muse cells.

**Figure 6.**
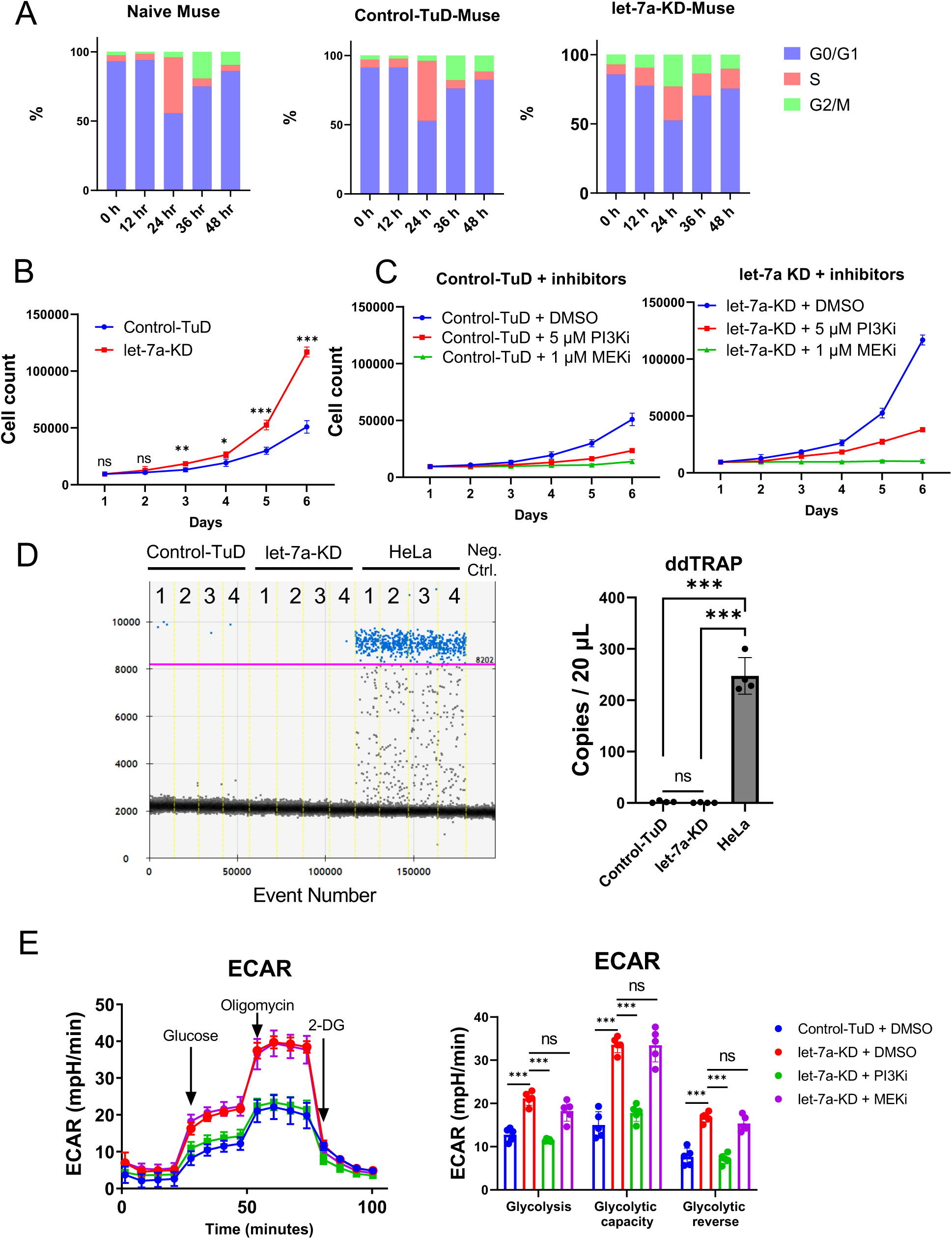
The influence of let-7 on cell proliferation and glycolysis. (A) Flow cytometry analysis of cell cycle variation among naïve, control-TuD-, and let-7a-KD-Muse cells by PI staining for 48 h. (B) Cell proliferation assay in control-TuD and let-7a-KD Muse cells by cell count (n=4). (C) Cell proliferation assay in control -TuD and let-7a-KD Muse cells under the presence of DMSO, 5 μM PI3Ki, and 1 μM MEKi by cell count (all n=4). (D) Comparison of telomerase activity by ddTRAP among control-TuD, let-7a-KD-Muse cells, and HeLa cells (n=4). HeLa cells were used as a positive control. No template control during the extension reaction was used as a negative control. Neg. ctrl.: negative control. The threshold was set at 8202 (purple line). (E) Comparison of ECAR among DMSO-treated control-TuD and let-7a-KD-Muse cells in the presence of DMSO, PI3Ki, and MEKi (n=5). PI3Ki: LY294002. MEKi: PD0325901. ECAR: extracellular acidification rate. 2-DG: 2-deoxy-D-glucose.

The effect of PI3Ki and MEKi on the proliferation of let-7a-KD-Muse cells was also examined. In both control-TuD- and let-7a-KD-Muse cells, PI3Ki suppressed proliferation activity compared with that of DMSO treated-Muse cells, while MEKi induced growth arrest in both types of Muse cells (Fig. 6C). Immunocytochemistry showed that MEKi treatment decreased Ki67 levels to under the detection limit in both control-TuD- and let-7a-KD-Muse cells after day 3, while PI3Ki treatment induced no difference in either Muse cell type compared with DMSO treated-Muse cells at day 5 (Supplemental Fig. S5A and S5B). These results demonstrated that PI3Ki suppressed proliferation activity, and MEKi induced cell cycle arrest in Muse cells.

Telomerase activity is one of the indicators that reflect the replicative immortality of cancer cells. (50). Droplet digital telomere repeat amplification protocol (ddTRAP) demonstrated that the telomerase activity was under the detection limit in both control-TuD and let-7a-KD Muse cells but was significantly higher in HeLa cells (Fig. 6D), suggesting that let-7a KD did not clearly evoke the activation of telomerase in Muse cells.

### Let-7 inhibited glycolysis through the PI3K-AKT pathway in Muse cells

To investigate the effect of let-7a on cellular metabolism in Muse cells, we measured the extracellular acidification rate (ECAR), an indicator of glycolysis. In let-7a-KD-Muse cells, glycolysis, glycolytic capacity, and glycolytic reversal were significantly increased compared with those in control-TuD-Muse cells (p<0.001) (Fig. 6E), suggesting that let-7 originally suppressed ECAR. PI3Ki treatment reversed the enhanced ECAR in let-7a KD-Muse-cells (Fig. 6E). Glycolysis, glycolytic capacity, and glycolytic reversal were all significantly suppressed by PI3Ki (p<0.001) compared with that in DMSO-treated let-7a-KD-Muse cells. At the same time, MEKi treatment did not have such an effect (Fig. 6E). Therefore, let-7 was suggested to inhibit glycolysis through the PI3K pathway, but not through the MEK/ERK pathway.

## Discussion

Muse cells are pluripotent-like endogenous somatic stem cells that are non-tumorigenic. In this study, we identified a unique mechanism by which Muse cells maintain pluripotency gene expression without being tumorigenic. In contrast to ESCs and iPSCs, whose pluripotency depends on LIN28, Muse cells basically do not express LIN28 and their pluripotency gene expression is sustained by the tumor suppressor miRNA let-7. LIN28 positively regulates pluripotency gene expression in PSCs (14, 51). Not only in ESCs and iPSCs, LIN28 is abundantly expressed in the early phase of development as represented by 2-cell stage embryos and blastocysts (13), but gradually decreases with differentiation/development and is eventually confined to a few somatic stem cells such as spermatogonial stem cells and neural stem cells in adults (52, 53). Recurrence of LIN28 expression in somatic cells is associated with some kinds of cancers (17, 54). Therefore, LIN28 is considered to be an oncogene. In the present study, the loss-of-function experiment demonstrated that let-7 could drive sustained expression of pluripotency genes by inhibiting the PI3K-AKT pathway in Muse cells (Fig. 5). A system utilizing the tumor suppressor miRNA let-7 rather than LIN28 to maintain pluripotent gene expression provides a low-risk model that enables both pluripotency and non-tumorigenicity. A summary of the proposed schema is shown in Fig. 7.

**Figure 7.**
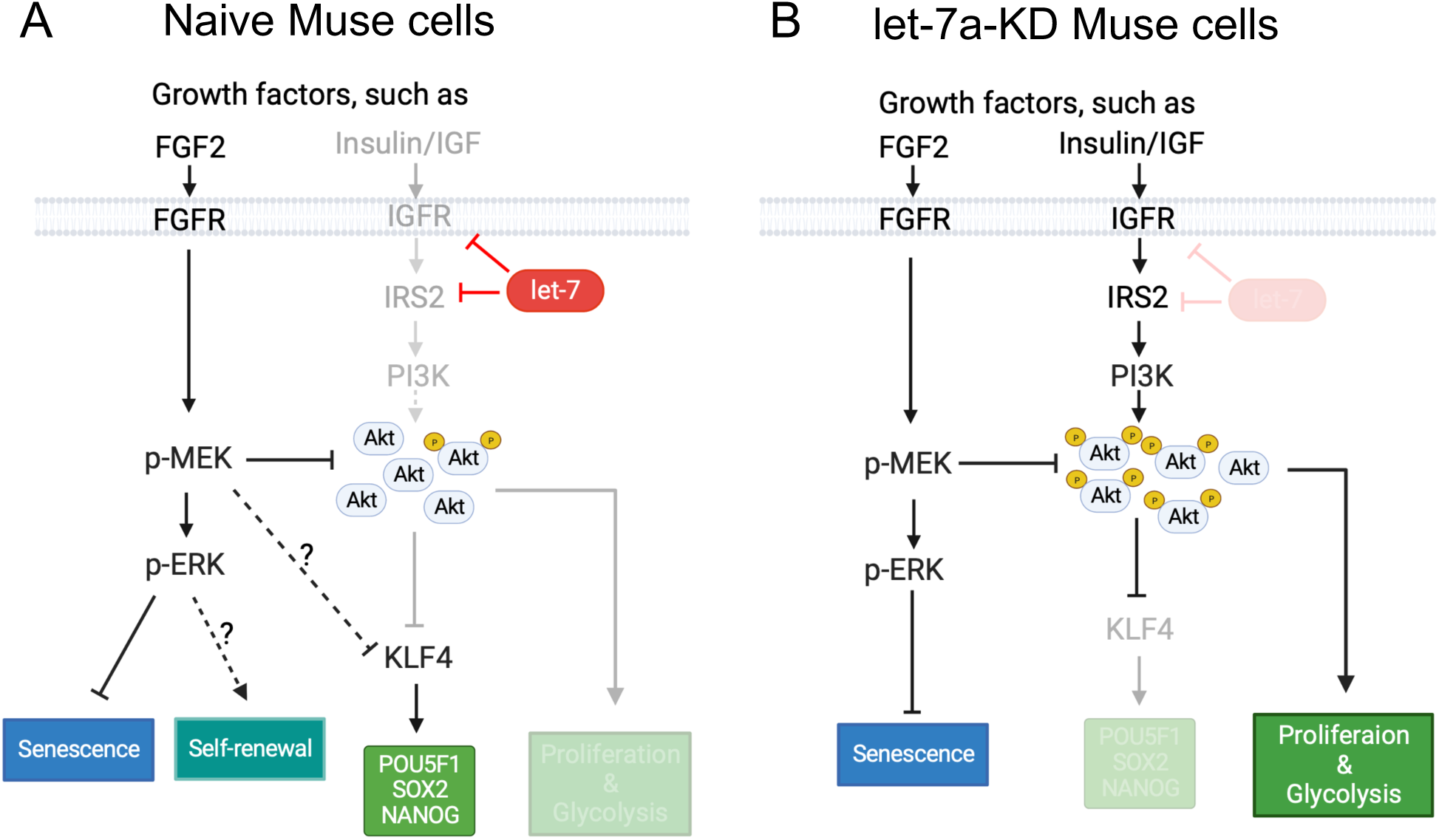
Summary. (A) The effect of let-7 on the PI3K-AKT and MEK/ERK pathways in naïve Muse cells. In naïve Muse cells, let-7 maintains the expression of pluripotency genes and inhibits proliferation and glycolysis. Let-7 inhibits the expression of IGF1R and IRS2 to repress the PI3K-AKT pathway. The PI3K-AKT pathway negatively controls the expression of *KLF4*, which promotes the expression of *POU5F1*, *SOX2*, and *NANOG*. The PI3K-AKT pathway directly inhibits the proliferation and glycolysis of naïve Muse cells. The other pathway, the MEK/ERK pathway, seems not affected by let-7. In naïve Muse cells, the MEK/ERK pathway suppresses the PI3K-AKT pathway by reducing the phosphorylation level of AKT. In addition, the MEK/ERK pathway also inhibits the expression of *KLF4*, but it virtually does not affect the expression of pluripotency genes. The MEK/ERK pathway is suggested to suppress senescence and maintain self-renewal of naïve Muse cells. (B) The effect of let-7 KD on the PI3K-AKT and MEK/ERK pathways in let-7-KD Muse cells. The PI3K-AKT pathway is more activated in let-7-KD-Muse cells than in naïve Muse cells. The increased AKT phosphorylation inhibits the expression of *KLF4*, which leads to the downregulation of the pluripotency genes *POU5F1*, *SOX2*, and *NANOG*. Cell proliferation and glycolysis were promoted. The MEK/ERK pathway reduces the phosphorylation level of AKT. The MEK/ERK pathway also suppresses cell senescence.

One outstanding difference between Muse cells and ESCs is the function of the PI3K-AKT pathway. In mouse ESCs, knockout of PTEN, a negative regulator of the PI3K-AKT pathway, upregulates KLF4, POU5F1, and NANOG levels (55). In human ESCs, PI3K inhibition leads to decreased pluripotency gene expression (56, 57). Thus, the PI3K-AKT pathway positively regulates pluripotency gene expression in ESCs. In Muse cells, however, we found that let-7 inhibited the expression of IGF1R and IRS2, leading to the suppression of PI3K-AKT downstream of IGF1R and IRS2, and sustained *KLF4* expression (Figs. 3–5). Consistently, PI3K-AKT inhibited the expression of *KLF4,* the factor upstream of *NANOG*, *POU5F1*, and *SOX2*; in naïve Muse cells*, KLF4, NANOG*, *POU5F1*, and *SOX2* were upregulated by PI3Ki treatment (Fig. 5C), and let-7a KD induced downregulation of these factors (Fig. 5D), which was counteracted by PI3Ki (Fig. 5F). These findings indicate that the PI3K-AKT pathway negatively regulates the expression of pluripotency genes in Muse cells, and that the inhibitory function of let-7 against PI3K-AKT is important for maintaining the expression of pluripotency genes.

We also found that KLF4 is a key regulator for sustaining pluripotency gene expression in Muse cells, as indicated by the suppression of *POU5F1*, *SOX2*, and *NANOG* after KLF4-siRNA introduction (Fig. 5H). In ESCs, KLF4 is downstream of the JAK/STAT3 pathway (48) and mediates the recruitment of Cohesin to the enhancer of *Pou5f1* to regulate *Pou5f1* transcription (58). Furthermore, KLF4 directly binds to the *Nanog* promoter to drive its expression (49). In this manner, KLF4 is able to act as the key regulator of pluripotency gene expression. Further studies are needed to determine how the PI3K-AKT pathway inhibits *KLF4* expression in Muse cells. The PI3K-AKT pathway is reported to activate the mammalian target of rapamycin complex 1 (mTORC1), leading to the inhibition of transcription factor E74-like factor 4, known to directly activate the expression of *KLF4* in T cells (59). A similar mechanism might be functioning in Muse cells, but more detailed studies are required to elucidate whether or not the PI3K-AKT-mTORC pathway mediates KLF4 expression.

Let-7 is suggested to participate in the suppression of proliferation and glycolysis in Muse cells through inhibiting the IGF1R-IRS2-PI3K-AKT pathway (Fig. 6). Some cell cycle-relevant factors, such as cyclin D1, cyclin D2, CDK4, CDK6, and cell division cycle 25A, are reported to be let-7 targets (17, 60, 61). Therefore, let-7 could suppress cell cycle activity by directly inhibiting those factors. In our microarray analysis, some of the cell cycle-relevant factors were upregulated in let-7 KD Muse cells (Fig. S2E), which might be the reason why let-7 KD accelerated proliferation. Glucose transporter type 4 increases glucose uptake under regulation of the PI3K-AKT pathway (62, 63). By inhibiting the PI3K-AKT pathway, let-7 may inhibit glycolysis indirectly. In T cells, let-7 suppresses glycolysis by directly targeting hexokinase 2 mRNA, a critical enzyme in glycolysis (64). Thus, it is possible that let-7 inhibits glycolysis in a direct or indirect manner in Muse cells. Further studies are needed to elucidate the detailed mechanisms. The telomerase activity was reported to be lower in Muse cells than that in HeLa cells and iPSCs (27, 29, 65). Let-7a KD stimulated the proliferation but did not increase the telomerase activity, suggesting that let-7 is a key factor for controlling cell cycle but is not directly connected to tumorigenic proliferation in Muse cells.

Overactivation of the PI3K-AKT pathway based on genetic alterations of factors such as PIK3CA, PTEN, and AKT is frequently observed in various types of tumors (66). After birth, the role of stem cells gradually changes from the rapid proliferation phase in the early stage of embryonic development to a moderate/slow proliferation phase in adult tissue where stem cells are required for tissue repair and cell replenishment (67). Inhibition of the PI3K-AKT pathway plays a pivotal role in maintaining the quiescent state of somatic stem cells, such as muscle stem cells and hematopoietic stem cells (68–70). High proliferation levels seem unfavorable for somatic stem cells to maintain a quiescent state. In ESCs, iPSCs, and cancer cells, high proliferation tends to accompany glycolytic metabolism, even in an oxygen-sufficient environment, to produce biomass (71). In Muse cells, let-7 may be a barrier that prevents cells from unlimited proliferative activity.

The MEK/ERK pathway is suggested to participate in controlling cell cycle and proliferative activity through supporting self-renewal and suppressing senescence in Muse cells (Fig. 6 and S5), while not being directly involved in pluripotency gene expression (Fig. 5). The MEK/ERK pathway exhibits opposite effects between naïve and primed ESCs: Inhibition of ERK facilitates maintenance of the naïve pluripotent state, while on the other hand, activation of the MEK/ERK pathway by FGF2 maintains the primed pluripotent state (Weinberger et al., 2016). In Muse cells, FGF2 withdrawal (Fig. 1A) and/or MEK/ERK pathway inhibition (Fig. 5A) decreased the percentage of Muse cells among BM-MSCs, suggesting that the FGF2-MEK/ERK pathway may play a role in maintaining self-renewal of Muse cells. The MEK/ERK pathway, however, seems to have a limited effect on the expression of pluripotency genes in Muse cells (Fig. 5C). We also analyzed the possibility that the MEK/ERK pathway inhibits senescence. A flattened morphology, increased SA-βgal activity, and arrested cell cycle are features of senescent cells (72). Both control-TuD and let-7-KD Muse cells exhibited a flattened morphology (Fig. 5B and 5E), arrested proliferation (Fig. 6C), and loss of the Ki67 expression (Fig. S5) when treated with a MEKi. MEKi-treated let-7-KD Muse cells showed higher SA-βgal activity than control-TuD Muse cells (Fig. S3A), implying that let-7 also plays a role in inhibiting cellular senescence when the MEK/ERK pathway is suppressed. The fact that let-7-KD Muse cells underwent senescence easier than control-TuD Muse cells might explain why pluripotency gene expression decreased sharply 1 day after MEKi treatment in let-7-KD Muse cells but not in naive Muse cells (Fig. 5C, F).

There are limitations to this study. Firstly, in this study, we found that MEK inhibited p-AKT (Fig. 4E), and p-AKT inhibited the expression of *KLF4* (Fig. 5C) in naïve Muse cells. Given these findings, inhibition of MEK was originally proposed to decrease the expression of *KLF4* via the activation of the PI3K-AKT pathway. However, our result showed that the inhibition of MEK upregulated the expression of *KLF4* in naïve Muse cells (Fig. 5C). Thus, MEK may regulate the expression of *KLF4* in Muse cells by an unknown PI3K-AKT-independent mechanism. The mechanism of how MEK regulates the expression of *KLF4* needs to be clarified in the future. Secondly, KLF4 was suggested to locate upstream of the three pluripotency genes by the KLF4 knockdown experiment (Fig. 5H). However, the upregulated *KLF4* expression induced by MEKi did not increase the expression of POU5F1, SOX2, and NANOG (Fig. 5C). This may be explained by the potential cell cycle arrest of Muse cells after MEKi treatment (Fig. 6C and Fig. S5) since cell cycle arrest was previously reported to decrease the expression of pluripotency genes (73). The detailed mechanism should be clarified in the future. Thirdly, Muse cells could overexpress pre-let-7a-3 but not mature let-7a. On the other hand, NTERA2 could overexpress both pre-let-7a-3 and mature let-7a. This suggested that the lentiviral overexpression system worked in NTERA2 but not in Muse cells. Muse cells may have a unique defense mechanism to prevent overexpression of let-7, which remains to be elucidated.

In the present study, we demonstrated a novel role of tumor suppressor miRNA let-7 as a key player for maintaining pluripotency gene expression by inhibiting the PI3K-AKT pathway, as well as the importance of MEK/ERK pathway in suppressing apoptosis and senescence in endogenous pluripotent-like non-tumorigenic Muse cells.

## Experimental procedures

### Cell culture

Human mesenchymal stem cells (hMSCs, LONZA, PT-2501), normal human dermal fibroblasts (NHDFs, LONZA, CC-2511), HEK293T, NTERA2, HeLa cells, and induced pluripotent stem cells (iPSCs) were used in this research. hMSCs and NHDFs were maintained in Minimum Essential Medium Eagle (αMEM, MilliporeSigma, M4526) supplemented with 10% fetal bovine serum (Hyclone, SH30910.03), 1x GlutaMAX (Gibco, Thermo Fisher Scientific, 35050-061), 1 ng/mL human basic FGF2 (Miltenyi Biotech, 130-093-840) and kanamycin (Gibco, 15160-054). Culture medium was exchanged every 2 days. FGF2 was kept at 4°C for no longer than 1 week and was freshly added while preparing the growth medium. HEK293T, NTERA2, and HeLa cells were maintained in Dulbecco’s modified Eagle’s medium (Gibco, 11965-092) supplemented with 10% fetal bovine serum, 1 mM sodium pyruvate (Gibco, 11360-070), and kanamycin. iPSCs were induced from NHDFs as previously described (29) and maintained in StemFit AK02N (AJINOMOTO, RCAK02N). The inhibitors used in this study were LY294002 (Selleck, S1105), PD0325901 (Wako, 162-25291), and MK2206 (Selleck, S1078). All the cells were cultured in a humidified incubator with 5% CO_2_ at 37°C.

### Muse cell sorting

Muse cells were sorted when hMSCs or NHDFs reached 100% confluency. For antibody labeling, the cells were incubated with anti-SSEA-3 rat IgM antibody (1:1000, BioLegend, 330302) at 4°C for 1 h following incubation with fluorescein (FITC) AffiniPure goat anti-rat IgM (1:100, Jackson ImmunoResearch, 112-095-075) or allophycocyanin-conjugated (APC) AffiniPure F (ab’)_2_ fragment goat anti-Rat IgM (1:100, Jackson ImmunoResearch, 112-136-075) at 4°C for 1 h. Purified Rat IgM, κ Isotype Ctrl Antibody (BioLegend, 400801) was used as a negative control for gate setting. FACS buffer (5% bovine serum albumin [BSA], 2 mM EDTA, and FluoroBrite Dulbecco’s modified Eagle’s medium [Thermo Fisher Scientific, A1896701]) was used for diluting the antibodies. Muse cells were collected using a BD FACSAria II SORP Flow Cytometer Cell Sorter (Becton Dickinson) in purify mode.

### Reverse transcription PCR (RT-PCR)

Total RNA was isolated with a mirVana™ miRNA Isolation Kit (Invitrogen, Thermo Fisher Scientific, AM1560) following the manufacturer’s instructions. The quality and concentration of the total RNA were measured with a Nanodrop 1000 spectrophotometer (Thermo Fisher Scientific). cDNAs were generated from mRNAs by RT-PCR with a SuperScript III first-strand synthesis system (Invitrogen, 18080044) and oligo(dT)20 primers (Invitrogen, 18418020) using a Takara Thermal PCR Cycler (Takara Bio). RT-PCR of miRNAs was performed with a Taqman microRNA Reverse Transcription Kit (Invitrogen, 4366597).

### Quantitative PCR (qPCR)

qPCR was performed with either Taqman Universal Master Mix II, using UNG (Applied Biosystems, 4440038) or PowerUP SYBR Green Master Mix (Applied Biosystems, A25742) running in the 7500 Fast Real-Time PCR System (Applied Biosystems). When performing SYBR Green qPCR assays, we confirmed the melting curve to ensure the specificity of the PCR products. ACTB and RNU48 were used for mRNA and miRNA qPCR as endogenous controls, respectively. The 2^-ΔΔCT^ relative quantification method was used in all analyses for calculation. Primers used in this study were obtained in 3 ways: primer BLAST service was provided by the National Center for Biotechnology Information (NCBI), ordering of Taqman qPCR probes, and primer searches on Primer Bank (https://pga.mgh.harvard.edu/primerbank/). Supplemental Table 1 shows the details of the primers.

### Droplet digital PCR (ddPCR)

#### Gene expression assay

Total RNA and cDNA were extracted and synthesized as described above. Taqman gene expression probes of LIN28A/B were used (Supplemental Table 1). cDNA samples and Droplet Generator Oil for Probe (Bio-Rad, 186-3005) were loaded in a DG8 cartridge (Bio-Rad, 186-4008), respectively. The QX200 Droplet Generator (Bio-Rad) was used to mix the samples and oil to generate the droplets. The mixed droplets were transferred to a 96-well twin.tec PCR Plate (Eppendorf, 95579) before sealing the plate with Foil Heat Seal (Bio-Rad, 1814040) in a PX1 PCR Plate Sealer (Bio-Rad). PCR was carried out on a C1000 Touch Thermal Cycler (Bio-Rad) at the following temperature and time: denaturation 95°C, 30s; annealing/extension 60°C, 1 min for 40 cycles. The results were read in a Droplet Reader (Bio-Rad) and analyzed by QuantaSoft (Bio-Rad).

#### DdTRAP

Telomerase activity was measured by ddTRAP following previously published protocol (74). A whole cell lysate was prepared by adding NP40 lysis buffer. Telomerase extension (TS) primer (5’-AATCCGTCGAGCAGAGTT) was used in the extension reaction (25°C, 1 h; 95°C, min; 12°C hold). In the extension reaction, no template control was used as a negative control. TS primer and ACX (reverse amplification) primer (5’-GCGCGGCTTACCCTTACCCTTACCCTAACC) were used to amplify the telomerase-extended substrates (95°C, 30 s; 54°C, 30s; 72°C, 30s for 40 cycles). Droplet Generation Oil of EvaGreen (Bio-Rad, 186-4006) was used. The results were read in a Droplet Reader (Bio-Rad) and analyzed by QuantaSoft (Bio-Rad).

### Western blotting

Cells were washed with 1x cold PBS twice and lysed by adding RIPA Lysis and Extraction Buffer (Thermo Fisher Scientific, 89900). A protease inhibitor (Thermo Fisher Scientific, 87785) and phosphatase inhibitor (Roche, 04906837001) were added to inhibit the protease and phosphatase activity. Cell lysates were centrifuged at 13,000 rpm at 4°C for 10 min. The supernatant was transferred to new tubes, and protein quantification was performed using BCA Protein Assay Kit (Thermo Fisher Scientific, 23225) following the manufacturer’s instructions. Sodium dodecyl sulfate-polyacrylamide gel electrophoresis (SDS-PAGE) was carried out with SDS-PAGE gels, and proteins were transferred to polyvinylidene difluoride membranes (Millipore, IPVH00010). Membranes were blocked in 5% skim milk (Nacalai, 31149-75)/Tris-buffered saline with 0.05% Tween-20 (TBST) for 1 h and incubated with primary antibodies overnight at 4°C. Secondary antibody reactions were performed at room temperature for 1 h. The blots were washed 3 times with TBST at room temperature for 5 min after the primary or secondary antibody reactions. The blots were developed by Pierce ECL Plus Western Blotting Substrate (Thermo Fisher Scientific, 32132). Images were acquired using Fusion FX imaging systems (Vilber). Band intensity was quantified by ImageJ software (75). The primary and secondary antibodies used in this research are listed in Supplemental Table 2.

The following antibody dilutions were used: LIN28A (1:1000, Cell Signaling), LIN28B (1:1000, Cell Singling), IGF1 receptor β (1:1000, Cell Signaling), Insulin receptor β (1:1000, Cell Signaling), NRAS (1:200, Santa Cruz Biotechnology), p-AKT (T308) (1:1000, Cell Signaling), p-AKT (S473; 1:1000, Cell Signaling), AKT (1:1000, Cell Signaling), Phospho-p44/p42 MEK/ERK (ERK1/2; Thr202/Tyr 204) (1:1000, Cell Signaling), p44/p42 MEK/ERK (ERK1/2; 1:1000, Cell Signaling), β-actin (1:10000, Abcam), and KLF4 (1:1000, Cell Signaling).

### let-7 knockdown

pWPXL was a gift from Professor Didier Trono (Addgene #12257). Each pWPXL-U6-TuD-EF1α-GFP (or mCherry) of let-7a, -7b, -7e, and -7i was constructed. Lentivirus was created by the transduction of each plasmid with Lipofectamine 3000 (Invitrogen, L3000075) into HEK293T cells. Lentivirus was collected and concentrated using Ambion Ultra-15 centrifugal filters (Millipore, UFC910024) 72 h after transduction. The hMSCs were transfected with each lentivirus for 3 days. To allow for adequate collection of Muse cells after lentivirus infection, hMSCs were also cultured for 3 to 4 PDLs. TuD-let-7-transfected hMSCs were then harvested and labeled with anti-SSEA-3 antibody and either FITC or APC secondary antibody for sorting using the BD FACSAria II SORP Flow Cytometer Cell Sorter (Becton Dickinson).

### Luciferase assay

pGL4.13 (Promega, E6681) and pRL-CMV (Promega, E226A) are commercially available. The plasmid was first cut by XbaI and HindIII. A restriction enzyme BssHII site was then added artificially. The let-7 complementary sequences and ath-mir-416 complementary sequence yielded by annealing specific oligo sets were inserted into the restriction enzyme sites BssHII and XbaI of pGL4.13 to obtain pFluc-let-7a, pFluc-let-7b, pFluc-let-7e, and pFluc-let-7i plasmids. Let-7a, -7b, 7e, and -7i KD hMSC-Muse cells were sorted and plated on 12-well plates at a density of 15,000 cells/cm^2^. The modified pGL4.13 plasmids and pRL-CMV plasmid were co-transfected with Lipofectamine 2000 (Invitrogen) at a mass ratio of 30:1. Naive, control-TuD, and each type of let-7-KD-Muse cell were all transfected by pFluc-let-7a, pFluc-let-7b, pFluc-let-7e, and pFluc-let-7i plasmids (Fig. S2B). Luciferase assay was performed after 24 h using the Dual-Luciferase Reporter Assay System (Promega, E1910).

### Microarray and microarray analysis

Naïve, TuD-control-, let-7a-KD-, let-7b-KD-, let-7e-, and let-7i-KD-Muse cells were collected from hMSCs. Total RNA extraction, cDNA synthesis and hybridization, and microarray analysis were completed by Takara Bio Inc. The SurePrint G3 Human GE v3 8×60K microarray (Agilent Technologies) was used for hybridization. Slides were scanned on the Agilent SureScan Microarray Scanner (G2600D) using the 1-color scan setting. The scanned images were analyzed by Feature Extraction Software (Agilent Technologies).

#### Normalization

For calculating the scaling factor, a trimmed mean probe intensity was settled by removing 2% of the lower and higher end of the probe intensities. Using the scaling factor, the normalized signal intensities were calculated.

#### Showing differential expression by heatmap

Genes whose expression was under detection limit were filtered in Microsoft Excel, and the heatmap of differential expression was produced by the pheatmap package of R (https://CRAN.R-project.org/package=pheatmap).

### Cell starvation and growth factor treatment

Muse cells were seeded at 15,000 cells/cm^2^. After attaching, medium was changed to serum-free αMEM, and cells were further cultured for 8 h. After the addition of 100 ng/mL insulin, 100 ng/mL IGF1, and 2 ng/mL FGF2 to the corresponding wells, the cells were treated for 15 min. Cells were then lysed, and the samples were analyzed by Western blotting.

### Flow cytometry analysis of the ratio of the Muse population

Cells were seeded at 1 million/10 cm dish. After adherence, medium was changed to culture medium with FGF2, without FGF2, containing 5 µM LY294002, and containing 1 μM PD0325901, respectively. After culturing for 2 PDLs, cells were then collected and stained with anti-SSEA-3 antibody as described above. The positive ratio of SSEA-3+ cells was analyzed by the CytoFLEX S Flow Cytometer (Beckman Coulter). Flow cytometry data were analyzed by Kaluza Analysis Software (Beckman Coulter).

### Analysis of pluripotency gene expression

Naive and let-7-KD Muse cells were each collected and seeded at 15,000 cells/cm^2^. After attaching, cells were treated with DMSO, 5 µM LY294002, or 1 µM PD0325901, respectively, for 24 h. Gene expression was analyzed by qPCR as described above.

### Let-7 overexpression

The let-7 overexpression plasmids are commercially available from System Biosciences (SBI). The catalog numbers of the plasmids used in this research are PMIRH000PA-1 (empty vector), PMIRHlet7a3PA-1 (let-7a overexpression), PMIRHlet7bPA-1 (let-7b overexpression), PMIRHlet7ePA-1 (let-7e overexpression), and PMIRHlet7iPA-1 (let-7i overexpression). Lentivirus was created by transduction of each plasmid with Lipofectamine 3000 (Invitrogen, L3000075) into HEK293T cells. Lentivirus was collected and concentrated by Ambion Ultra-15 centrifugal filters (Millipore, UFC910024) 72 h after transduction. The hMSCs were transfected by each lentivirus for 3 days. To obtain enough number of Muse cells after lentivirus infection, hMSCs were also cultured for 3 to 4 PDLs. TuD-let-7-transfected hMSCs were then harvested and labeled with anti-SSEA-3 antibody and APC secondary antibody for sorting using the BD FACSAria II SORP Flow Cytometer Cell Sorter (Becton Dickinson).

### siRNA knockdown

Three kinds of Klf4 siRNA (Ambion, Thermo Fisher Scientific, ID: S17793, S17794, and S17795) were mixed up, and naive Muse cells were sorted and transfected with 25 pmol KLF4 siRNA or 25 pmol scrambled siRNA (Ambion) by Lipofectamine RNAiMAX (Invitrogen, 13778-150) in 6-well plates for 48 h following the manufacturer’s instructions. Total RNA was extracted, and gene expression was analyzed by qPCR as described above.

### Cell cycle analysis

Naive, control-TuD, let-7-KD-Muse cells were collected, respectively. 70% ethanol was pre-cooled at -30°C. Fix 120,000 cells from each group just after sorting with pre-cooled 70% ethanol at 4 °C for 30 minutes. The left cells were seeded as 25,000 cells/cm^2^ in 12 wells. Cells were detached from the culture plates every 12 hours and then fixed with pre-cooled 70% ethanol. After fixation, all the samples were stored at 4 °C. Centrifuge all the samples at 1000xg for 5 minutes at room temperature and wash with FACS buffer. Centrifuge again at 1000xg for 5 minutes at room temperature. Digest cells with 250 µg/mL RNase at 37°C for 15 minutes. Add Propidium Iodide (PI) to each sample and incubate at 4 °C for 10 minutes. Analyze the samples by CytoFLEX S Flow Cytometer (Beckman Coulter). All the data were analyzed by Kaluza Analysis Software (Beckman Coulter).

### Cell proliferation assay

Cells were seeded in 4-well plates at 5000 cells/well. Cells were trypsinized and counted by hemacytometers (WakenBtech, WC2-100) every 24 h until day 6.

### ECAR analysis

We seeded 10,000 cells per well in Seahorse XF96 cell culture microplates (Agilent Technologies, 103725-100) and pre-cultured cells 12 h before measurement. ECAR was then measured using the Seahorse XFe96 Analyzer (Agilent Technologies). We calculated fundamental parameters such as glycolysis, glycolytic capacity, and glycolytic reverse of ECAR following the manufacturer’s instructions.

### Flow cytometry analysis of apoptosis

#### TUNEL assay

Muse cells were collected and fixed with fresh 4% paraformaldehyde in PBS, pH 7.4 for 30 min, further incubated with permeabilization solution (0.1% Triton x100 in 0.1% sodium citrate solution) for 2 min on ice, and labeled by the DeadEnd Fluorometric TUNEL System (Promega, G3250) following the manufacturer’s instructions. A positive control was prepared with a DNase-digested sample. Apoptosis of Muse cells was observed by flow cytometry using a BD FACSAria II SORP Flow Cytometer Cell Sorter.

#### Annexin V staining

hMSCs treated with UV for 5 min followed by incubation for an additional 24 h were used as a positive control. Muse cells were seeded 20,000 cells/cm^2^ in a 24-well-plate and the medium was changed to DMSO with the addition of 5 μM LY294002 or 1 μM PD0325901 after cell attachment. Cells were collected 1, 3, and 5 days after inhibitor treatment and stained using a MEBCYTO Apoptosis Kit (Annexin V-FITC Kit; MBL, 4700) following the manufacturer’s instruction. Samples were analyzed by the CytoFLEX S Flow Cytometer (Beckman Coulter). Flow cytometry data were analyzed using Kaluza Analysis Software (Beckman Coulter).

### Flow cytometry analysis for senescence

Cells were stained with the SPiDER-βGal detection kit (Dojindo, SG02) following the manufacturer’s protocol and then collected by trypsinization. The mean fluorescence intensity (MFI) was analyzed by CytoFLEX S Flow cytometer (Beckman Coulter). The calculation method is shown below:

Sample A (experimental group): Cells stained with SPiDER-βGal

Sample B (experimental group): Cells without SPiDER-βGal staining

Sample C (control group): Cells stained with SPiDER-βGal

Sample D (control group): Cells without SPiDER-βGal staining

Variation of SA-β-gal activity = (MFI of A – MFI of B) – (MFI of C – MFI of D)

### Immunofluorescence

Cells were washed once with 1x PBS and fixed with 4% PFA/PBS for 30 min at 4 °C. The fixed cells were washed 3 times with 1x PBS and blocked with blocking solution for 30 min at room temperature before incubation with the primary antibody overnight at 4 °C. The next day, cells were washed 3 times with 1X PBS and incubated with the secondary antibody at room temperature for 30 min. Cells were then washed 3 times with 1x PBS and stained with DAPI for 5 min. The staining solution was discarded and changed to Milli-Q. The stained cells were observed and imaged with a fluorescence microscope (Keyence, BZ-X710).

Solutions and antibody dilutions were as follows: blocking solution (20% Blockace (KAC, UKB40), 5% BSA, 0.3% Triton X-100 in PBS); primary antibody dilution buffer (5% Blockace, 1% BSA, 0.3% Triton X-100 in PBS); secondary antibody dilution buffer (0.2% Triton X-100 in PBS). Ki67 (1:500, Abcam), and Alexa Fluor 488 AffiniPure F(ab’)□ Fragment Donkey Anti-Rabbit IgG (H+L) (1:200, Jackson ImmunoResearch).

### Statistical analysis

All statistical analyses were performed in Microsoft Excel or Graphpad Prism 8.0. Bioinformatic analysis was performed in R programming (Version 3.5.1). Data are presented as mean ± SD. The statistical significance of differences between 2 groups was calculated by the unpaired Student’s t-test. For comparison of more than 2 groups, 1-way ANOVA was conducted with Tukey’s post-hoc test. (*p < 0.05, **p < 0.01, ***p < 0.001, ns: no significance).

### Data Availability

All data are contained in the article and supporting information.

## Supporting Information

This article contains supporting information.

## Acknowledgments

We thank Minaka Sato for her technical support for cell culture and cell sorting. We thank Yoshihiro Kushida for his assistance in ddTRAP. We thank all the members of the Dezawa laboratory for their helpful discussions.

## Funding and additional information

This study was supported by Grant-in-Aid for Scientific Research B (20H04510) and Grant-in-Aid for Exploratory Research (19K22648) from the Ministry of Education, Science, and Culture of Japan, and collaborative research development with Life Science Institute, Inc.

## Author contributions

**Gen Li**: Methodology; conceptualization; investigation; data curation; formal analysis; investigation; writing - original draft; writing – review and editing. **Shohei Wakao**: Methodology. **Masaaki Kitada**: Methodology; conceptualization; investigation; supervision; writing - review and editing. **Mari Dezawa**: Methodology; conceptualization; funding acquisition; investigation; supervision; project administration; writing - review and editing.

## Reference

1. Ambros V, Horvitz HR. Heterochronic mutants of the nematode Caenorhabditis elegans. Science. 1984;226(4673):409–16.

2. Reinhart BJ, Slack FJ, Basson M, Pasquinelli AE, Bettinger JC, Rougvie AE, et al. The 21-nucleotide let-7 RNA regulates developmental timing in Caenorhabditis elegans. Nature. 2000;403(6772):901–6.

3. Lagos-Quintana M, Rauhut R, Meyer J, Borkhardt A, Tuschl T. New microRNAs from mouse and human. RNA. 2003;9(2):175–9.

4. Moss EG, Tang L. Conservation of the heterochronic regulator Lin-28, its developmental expression and microRNA complementary sites. Dev Biol. 2003;258(2):432–42.

5. Pasquinelli AE, McCoy A, Jimenez E, Salo E, Ruvkun G, Martindale MQ, et al. Expression of the 22 nucleotide let-7 heterochronic RNA throughout the Metazoa: a role in life history evolution? Evolution and Development. 2003;5(4):372–8.

6. Hagan JP, Piskounova E, Gregory RI. Lin28 recruits the TUTase Zcchc11 to inhibit let-7 maturation in mouse embryonic stem cells. Nature Structural & Molecular Biology. 2009;16(10):1021–5.

7. Heo I, Joo C, Cho J, Ha M, Han J, Kim VN. Lin28 Mediates the Terminal Uridylation of let-7 Precursor MicroRNA. Molecular Cell. 2008;32(2):276–84.

8. Bussing I, Slack FJ, Grosshans H. let-7 microRNAs in development, stem cells and cancer. Trends Mol Med. 2008;14(9):400–9.

9. Roush S, Slack FJ. The let-7 family of microRNAs. Trends Cell Biol. 2008;18(10):505–16.

10. Thornton JE, Gregory RI. How does Lin28 let-7 control development and disease? Trends Cell Biol. 2012;22(9):474–82.

11. Zhang J, Ratanasirintrawoot S, Chandrasekaran S, Wu Z, Scott, Yu C, et al. LIN28 Regulates Stem Cell Metabolism and Conversion to Primed Pluripotency. Cell Stem Cell. 2016;19(1):66–80.

12. Tsialikas J, Romer-Seibert J. LIN28: roles and regulation in development and beyond. Development. 2015;142(14):2397–404.

13. Vogt EJ, Meglicki M, Hartung KI, Borsuk E, Behr R. Importance of the pluripotency factor LIN28 in the mammalian nucleolus during early embryonic development. Development. 2012;139(24):4514–23.

14. Yu J, Vodyanik MA, Smuga-Otto K, Antosiewicz-Bourget J, Frane JL, Tian S, et al. Induced pluripotent stem cell lines derived from human somatic cells. Science. 2007;318(5858):1917–20.

15. Sekine K, Tsuzuki S, Yasui R, Kobayashi T, Ikeda K, Hamada Y, et al. Robust detection of undifferentiated iPSC among differentiated cells. Sci Rep-Uk. 2020;10(1).

16. Richards M, Tan SP, Tan JH, Chan WK, Bongso A. The Transcriptome Profile of Human Embryonic Stem Cells as Defined by SAGE. Stem Cells. 2004;22(1):51–64.

17. Balzeau J, Menezes MR, Cao S, Hagan JP. The LIN28/let-7 Pathway in Cancer. Front Genet. 2017;8:31.

18. Lv K, Liu L, Wang L, Yu J, Liu X, Cheng Y, et al. Lin28 Mediates Paclitaxel Resistance by Modulating p21, Rb and Let-7a miRNA in Breast Cancer Cells. PLoS ONE. 2012;7(7):e40008.

19. Wang L, Yuan C, Lv K, Xie S, Fu P, Liu X, et al. Lin28 Mediates Radiation Resistance of Breast Cancer Cells via Regulation of Caspase, H2A.X and Let-7 Signaling. PLoS ONE. 2013;8(6):e67373.

20. Yang X, Lin X, Zhong X, Kaur S, Li N, Liang S, et al. Double-Negative Feedback Loop between Reprogramming Factor LIN28 and microRNA let-7 Regulates Aldehyde Dehydrogenase 1–Positive Cancer Stem Cells. Cancer Research. 2010;70(22):9463–72.

21. Akao Y, Nakagawa Y, Naoe T. let-7 microRNA functions as a potential growth suppressor in human colon cancer cells. Biol Pharm Bull. 2006;29(5):903–6.

22. Liu C, Kelnar K, Vlassov AV, Brown D, Wang J, Tang DG. Distinct microRNA expression profiles in prostate cancer stem/progenitor cells and tumor-suppressive functions of let-7. Cancer Res. 2012;72(13):3393–404.

23. Takamizawa J, Konishi H, Yanagisawa K, Tomida S, Osada H, Endoh H, et al. Reduced expression of the let-7 microRNAs in human lung cancers in association with shortened postoperative survival. Cancer Res. 2004;64(11):3753–6.

24. Kumar MS, Erkeland SJ, Pester RE, Chen CY, Ebert MS, Sharp PA, et al. Suppression of non-small cell lung tumor development by the let-7 microRNA family. Proc Natl Acad Sci U S A. 2008;105(10):3903–8.

25. Aprile D, Alessio N, Demirsoy IH, Squillaro T, Peluso G, Di Bernardo G, et al. MUSE Stem Cells Can Be Isolated from Stromal Compartment of Mouse Bone Marrow, Adipose Tissue, and Ear Connective Tissue: A Comparative Study of Their In Vitro Properties. Cells. 2021;10(4):761.

26. Kuroda Y, Kitada M, Wakao S, Nishikawa K, Tanimura Y, Makinoshima H, et al. Unique multipotent cells in adult human mesenchymal cell populations. Proceedings of the National Academy of Sciences. 2010;107(19):8639–43.

27. Ogura F, Wakao S, Kuroda Y, Tsuchiyama K, Bagheri M, Heneidi S, et al. Human adipose tissue possesses a unique population of pluripotent stem cells with nontumorigenic and low telomerase activities: potential implications in regenerative medicine. Stem Cells Dev. 2014;23(7):717–28.

28. Sato T, Wakao S, Kushida Y, Tatsumi K, Kitada M, Abe T, et al. A Novel Type of Stem Cells Double-Positive for SSEA-3 and CD45 in Human Peripheral Blood. Cell Transplantation. 2020;29:096368972092357.

29. Wakao S, Kitada M, Kuroda Y, Shigemoto T, Matsuse D, Akashi H, et al. Multilineage-differentiating stress-enduring (Muse) cells are a primary source of induced pluripotent stem cells in human fibroblasts. Proceedings of the National Academy of Sciences. 2011;108(24):9875–80.

30. Alessio N, Squillaro T, Özcan S, Di Bernardo G, Venditti M, Melone M, et al. Stress and stem cells: adult Muse cells tolerate extensive genotoxic stimuli better than mesenchymal stromal cells. Oncotarget. 2018;9(27):19328–41.

31. Acar MB, Aprile D, Ayaz-Guner S, Guner H, Tez C, Di Bernardo G, et al. Why Do Muse Stem Cells Present an Enduring Stress Capacity? Hints from a Comparative Proteome Analysis. International Journal of Molecular Sciences. 2021;22(4):2064.

32. Uchida H, Niizuma K, Kushida Y, Wakao S, Tominaga T, Borlongan CV, et al. Human Muse Cells Reconstruct Neuronal Circuitry in Subacute Lacunar Stroke Model. Stroke. 2017;48(2):428–35.

33. Uchida N, Kushida Y, Kitada M, Wakao S, Kumagai N, Kuroda Y, et al. Beneficial Effects of Systemically Administered Human Muse Cells in Adriamycin Nephropathy. Journal of the American Society of Nephrology. 2017;28(10):2946–60.

34. Iseki M, Kushida Y, Wakao S, Akimoto T, Mizuma M, Motoi F, et al. Human Muse Cells, Nontumorigenic Phiripotent-Like Stem Cells, Have Liver Regeneration Capacity through Specific Homing and Cell Replacement in a Mouse Model of Liver Fibrosis. Cell Transplantation. 2017;26(5):821–40.

35. Yamada Y, Wakao S, Kushida Y, Minatoguchi S, Mikami A, Higashi K, et al. S1P–S1PR2 Axis Mediates Homing of Muse Cells Into Damaged Heart for Long-Lasting Tissue Repair and Functional Recovery After Acute Myocardial Infarction. Circulation Research. 2018;122(8):1069–83.

36. Wakao S, Oguma Y, Kushida Y, Kuroda Y, Tatsumi K, Dezawa M. Phagocytosing differentiated cell-fragments is a novel mechanism for controlling somatic stem cell differentiation within a short time frame. Cellular and Molecular Life Sciences. 2022;79(11).

37. Fujita Y, Nohara T, Takashima S, Natsuga K, Adachi M, Yoshida K, et al. Intravenous allogeneic multilineage-differentiating stress-enduring cells in adults with dystrophic epidermolysis bullosa: a phase 1/2 open-label study. Journal of the European Academy of Dermatology and Venereology. 2021;35(8).

38. Noda T, Nishigaki K, Minatoguchi S. Safety and Efficacy of Human Muse Cell-Based Product for Acute Myocardial Infarction in a First-in-Human Trial. Circulation Journal. 2020;84(7):1189–92.

39. Kuroda Y, Wakao S, Kitada M, Murakami T, Nojima M, Dezawa M. Isolation, culture and evaluation of multilineage-differentiating stress-enduring (Muse) cells. Nature Protocols. 2013;8(7):1391–415.

40. Haraguchi T, Ozaki Y, Iba H. Vectors expressing efficient RNA decoys achieve the long-term suppression of specific microRNA activity in mammalian cells. Nucleic Acids Res. 2009;37(6):e43.

41. Szklarczyk D, Morris JH, Cook H, Kuhn M, Wyder S, Simonovic M, et al. The STRING database in 2017: quality-controlled protein–protein association networks, made broadly accessible. Nucleic Acids Research. 2017;45(D1):D362–D8.

42. Manning BD, Toker A. AKT/PKB Signaling: Navigating the Network. Cell. 2017;169(3):381–405.

43. Gao L, Wang X, Wang X, Zhang L, Qiang C, Chang S, et al. IGF-1R, a target of let-7b, mediates crosstalk between IRS-2/Akt and MAPK pathways to promote proliferation of oral squamous cell carcinoma. Oncotarget. 2014;5(9):2562–74.

44. Zhu H, Shyh-Chang N, Segre AV, Shinoda G, Shah SP, Einhorn WS, et al. The Lin28/let-7 axis regulates glucose metabolism. Cell. 2011;147(1):81–94.

45. Kuppusamy KT, Jones DC, Sperber H, Madan A, Fischer KA, Rodriguez ML, et al. Let-7 family of microRNA is required for maturation and adult-like metabolism in stem cell-derived cardiomyocytes. Proc Natl Acad Sci U S A. 2015;112(21):E2785–94.

46. Melton C, Judson RL, Blelloch R. Opposing microRNA families regulate self-renewal in mouse embryonic stem cells. Nature. 2010;463(7281):621–6.

47. Jiang J, Chan YS, Loh YH, Cai J, Tong GQ, Lim CA, et al. A core Klf circuitry regulates self-renewal of embryonic stem cells. Nat Cell Biol. 2008;10(3):353–60.

48. Niwa H, Ogawa K, Shimosato D, Adachi K. A parallel circuit of LIF signalling pathways maintains pluripotency of mouse ES cells. Nature. 2009;460(7251):118–22.

49. Zhang P, Andrianakos R, Yang Y, Liu C, Lu W. Kruppel-like Factor 4 (Klf4) Prevents Embryonic Stem (ES) Cell Differentiation by Regulating Nanog Gene Expression. Journal of Biological Chemistry. 2010;285(12):9180–9.

50. Hanahan D, Robert. Hallmarks of Cancer: The Next Generation. Cell. 2011;144(5):646–74.

51. Hanna J, Saha K, Pando B, van Zon J, Lengner CJ, Creyghton MP, et al. Direct cell reprogramming is a stochastic process amenable to acceleration. Nature. 2009;462(7273):595–601.

52. Aeckerle N, Eildermann K, Drummer C, Ehmcke J, Schweyer S, Lerchl A, et al. The pluripotency factor LIN28 in monkey and human testes: a marker for spermatogonial stem cells? Mol Hum Reprod. 2012;18(10):477–88.

53. Cimadamore F, Amador-Arjona A, Chen C, Huang C-T, Terskikh AV. SOX2–LIN28/let-7 pathway regulates proliferation and neurogenesis in neural precursors. Proceedings of the National Academy of Sciences. 2013;110(32):E3017–E26.

54. Xiong H, Zhao W, Wang J, Seifer BJ, Ye C, Chen Y, et al. Oncogenic mechanisms of Lin28 in breast cancer: new functions and therapeutic opportunities. Oncotarget. 2017;8(15):25721–35.

55. Wang W, Lu G, Su X, Tang C, Li H, Xiong Z, et al. Pten-mediated Gsk3β modulates the naïve pluripotency maintenance in embryonic stem cells. Cell Death & Disease. 2020;11(2).

56. Armstrong L, Hughes O, Yung S, Hyslop L, Stewart R, Wappler I, et al. The role of PI3K/AKT, MAPK/ERK and NFκβ signalling in the maintenance of human embryonic stem cell pluripotency and viability highlighted by transcriptional profiling and functional analysis. Human Molecular Genetics. 2006;15(11):1894–913.

57. McLean AB, D’Amour KA, Jones KL, Krishnamoorthy M, Kulik MJ, Reynolds DM, et al. Activin A Efficiently Specifies Definitive Endoderm from Human Embryonic Stem Cells Only When Phosphatidylinositol 3-Kinase Signaling Is Suppressed. Stem Cells. 2007;25(1):29–38.

58. Wei Z, Gao F, Kim S, Yang H, Lyu J, An W, et al. Klf4 Organizes Long-Range Chromosomal Interactions with the Oct4 Locus in Reprogramming and Pluripotency. Cell Stem Cell. 2013;13(1):36–47.

59. Yamada T, Gierach K, Lee P-H, Wang X, Lacorazza HD. Cutting Edge: Expression of the Transcription Factor E74-Like Factor 4 Is Regulated by the Mammalian Target of Rapamycin Pathway in CD8+ T Cells. The Journal of Immunology. 2010;185(7):3824–8.

60. Johnson CD, Esquela-Kerscher A, Stefani G, Byrom M, Kelnar K, Ovcharenko D, et al. The let-7 microRNA represses cell proliferation pathways in human cells. Cancer Res. 2007;67(16):7713–22.

61. Schultz J, Lorenz P, Gross G, Ibrahim S, Kunz M. MicroRNA let-7b targets important cell cycle molecules in malignant melanoma cells and interferes with anchorage-independent growth. Cell Research. 2008;18(5):549–57.

62. Bryant NJ, Govers R, James DE. Regulated transport of the glucose transporter GLUT4. Nature Reviews Molecular Cell Biology. 2002;3(4):267–77.

63. Sano H, Eguez L, Teruel MN, Fukuda M, Chuang TD, Chavez JA, et al. Rab10, a Target of the AS160 Rab GAP, Is Required for Insulin-Stimulated Translocation of GLUT4 to the Adipocyte Plasma Membrane. Cell Metabolism. 2007;5(4):293–303.

64. Jiang S, Yan W, Wang SE, Baltimore D. Let-7 Suppresses B Cell Activation through Restricting the Availability of Necessary Nutrients. Cell Metab. 2018;27(2):393–403 e4.

65. Ogawa E, Oguma Y, Kushida Y, Wakao S, Okawa K, Dezawa M. Naïve pluripotent-like characteristics of non-tumorigenic Muse cells isolated from human amniotic membrane. Sci Rep-Uk. 2022;12(1).

66. Peng Y, Wang Y, Zhou C, Mei W, Zeng C. PI3K/Akt/mTOR Pathway and Its Role in Cancer Therapeutics: Are We Making Headway? Front Oncol. 2022;12:819128.

67. Zakrzewski W, Dobrzyński M, Szymonowicz M, Rybak Z. Stem cells: past, present, and future. Stem Cell Research & Therapy. 2019;10(1).

68. Elich M, Sauer K. Regulation of Hematopoietic Cell Development and Function Through Phosphoinositides. Front Immunol. 2018;9:931.

69. Gudmundsson KO, Du Y. Quiescence regulation by normal haematopoietic stem cells and leukaemia stem cells. FEBS J. 2022.

70. Wang G, Zhu H, Situ C, Han L, Yu Y, Cheung TH, et al. p110alpha of PI3K is necessary and sufficient for quiescence exit in adult muscle satellite cells. EMBO J. 2018;37(8):e98239.

71. Ito K, Suda T. Metabolic requirements for the maintenance of self-renewing stem cells. Nat Rev Mol Cell Biol. 2014;15(4):243–56.

72. Hernandez-Segura A, Nehme J, Demaria M. Hallmarks of Cellular Senescence. Trends Cell Biol. 2018;28(6):436–53.

73. Kareta MS, Sage J, Wernig M. Crosstalk between stem cell and cell cycle machineries. Current Opinion in Cell Biology. 2015;37:68–74.

74. Ludlow AT, Shelton D, Wright WE, Shay JW. ddTRAP: A Method for Sensitive and Precise Quantification of Telomerase Activity. Springer New York; 2018. p. 513–29.

75. Schneider CA, Rasband WS, Eliceiri KW. NIH Image to ImageJ: 25 years of image analysis. Nat Methods. 2012;9(7):671–5.

