## Supplementary figures and tables for "Tumor suppressor let-7 acts as a key regulator for maintaining pluripotency gene expression in Muse cells"

Li_Supplemental_Fig_S1

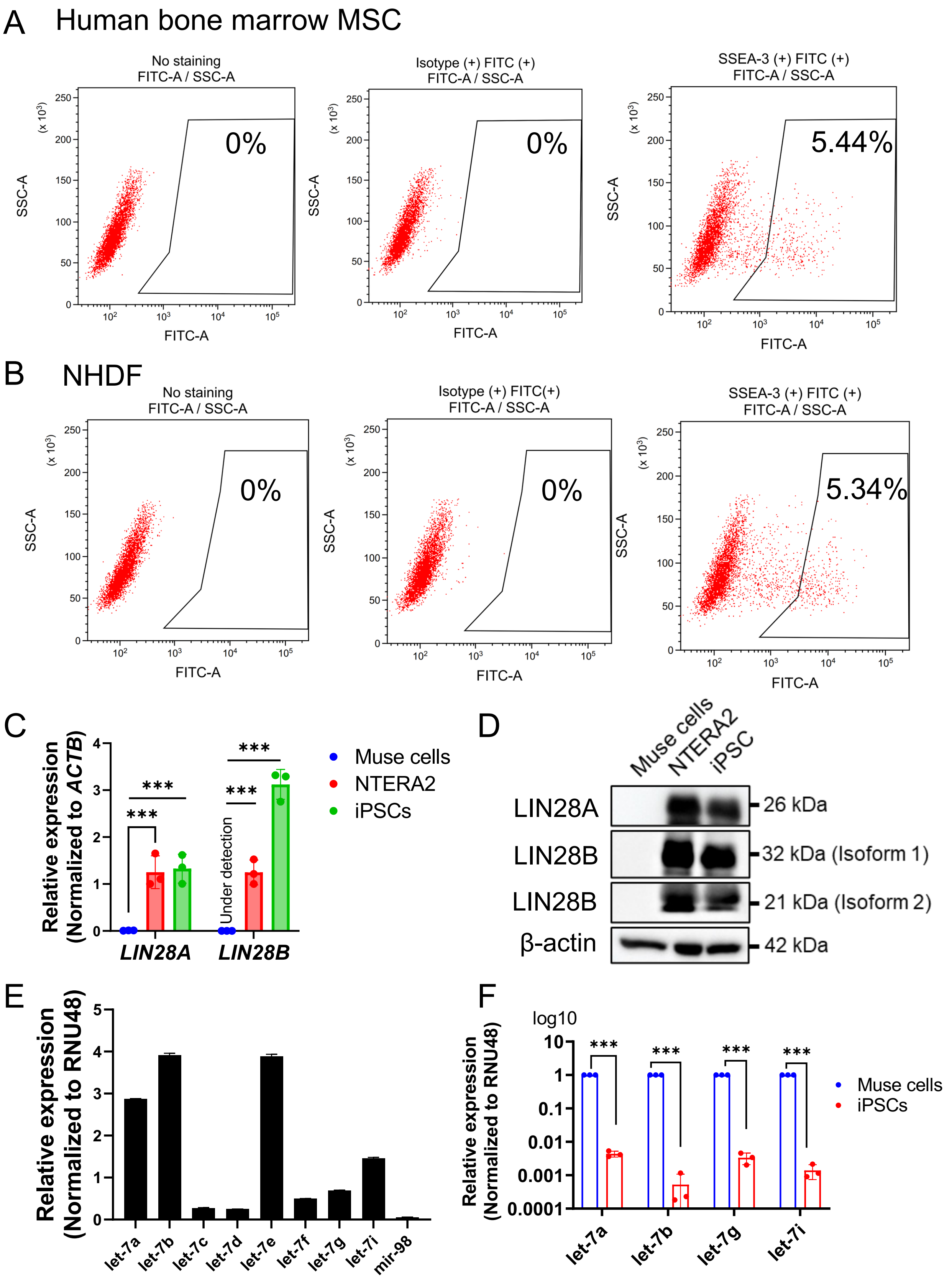

**Supplemental Figure S1. NHDF-Muse cells express let-7 but not LIN28.**

(A-B) FACS sorting of Muse cells.

(A) qPCR showing the expression of *LIN28A/B* in NHDF-Muse cells. ACTB was used as an endogenous control.

(B) Western blot showing LIN28A/B levels in NHDF-Muse cells, NTERA2, and iPSCs. β-Actin was used as an endogenous control.

(C) Expression of let-7 subtypes in NHDF-Muse cells (all n=3). RNU48 was used as an endogenous control.

(D) qPCR showing the expression of let-7a, -7b, -7e, and -7i in Muse cells and iPSCs (n=3). RNU48 was used as an endogenous control. A log10 scale was used for the y-axis.

### Li_Supplemental_Fig_S2

let-7 knockdown

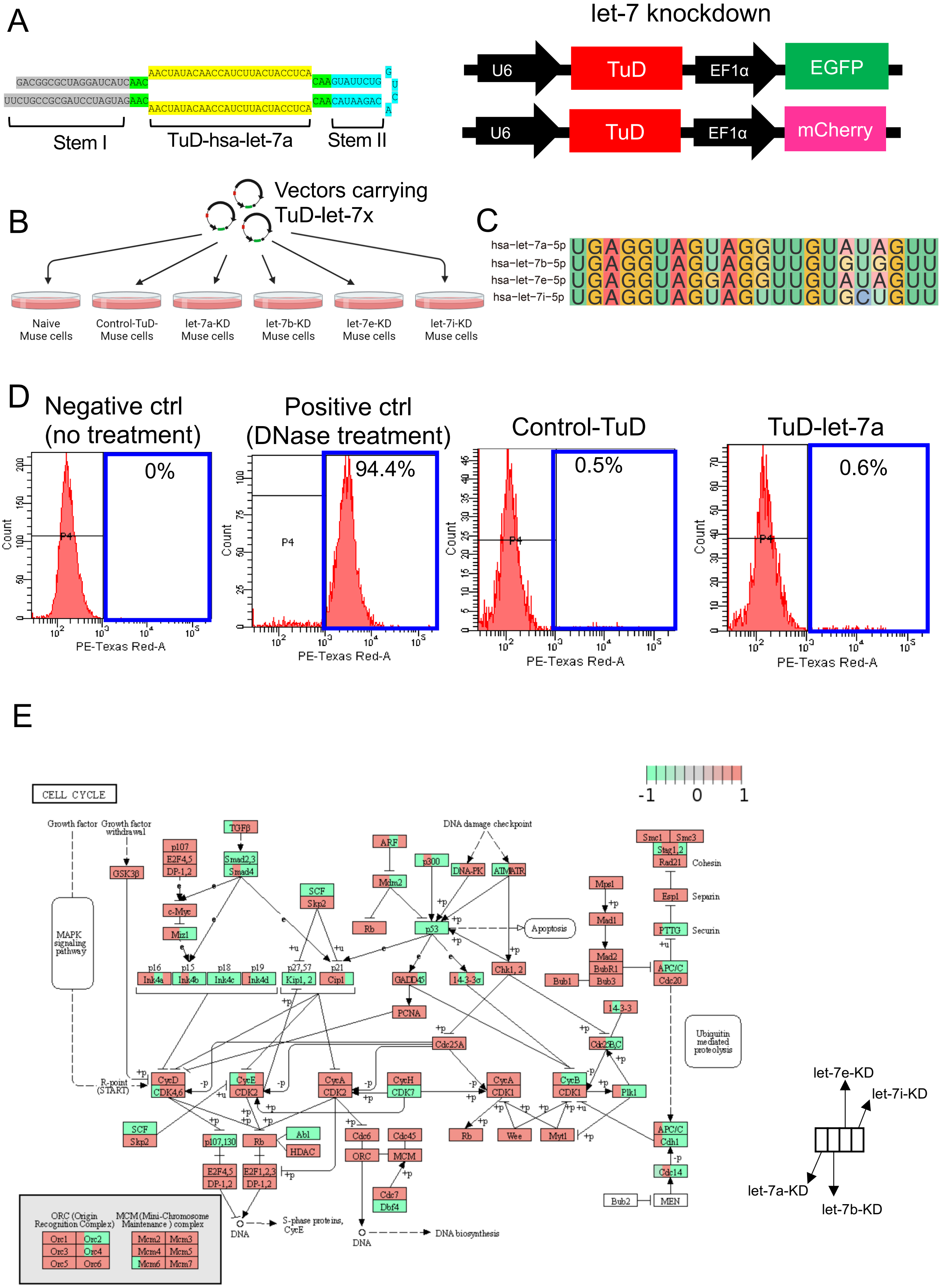

**Supplemental Figure S2. Let-7 knockdown.**

(A) Structure of TuD-hsa-let-7a (left) and construction of the lentiviral vector for let-7 knockdown (right).

(B) Scheme of the luciferase assay to check the knockdown effect of TuD-let-7. x in let-7x represents a, b, e, or i.

(C) Sequence comparisons of hsa-let-7a-5p, hsa-let-7b-5p, hsa-let-7e-5p, and hsa-let-7i-5p.

(D) Examination of cellular apoptosis by TUNEL assay. Ctrl: control

(E) KEGG pathway analysis of the cell cycle.

Li_Supplemental_Fig_S3

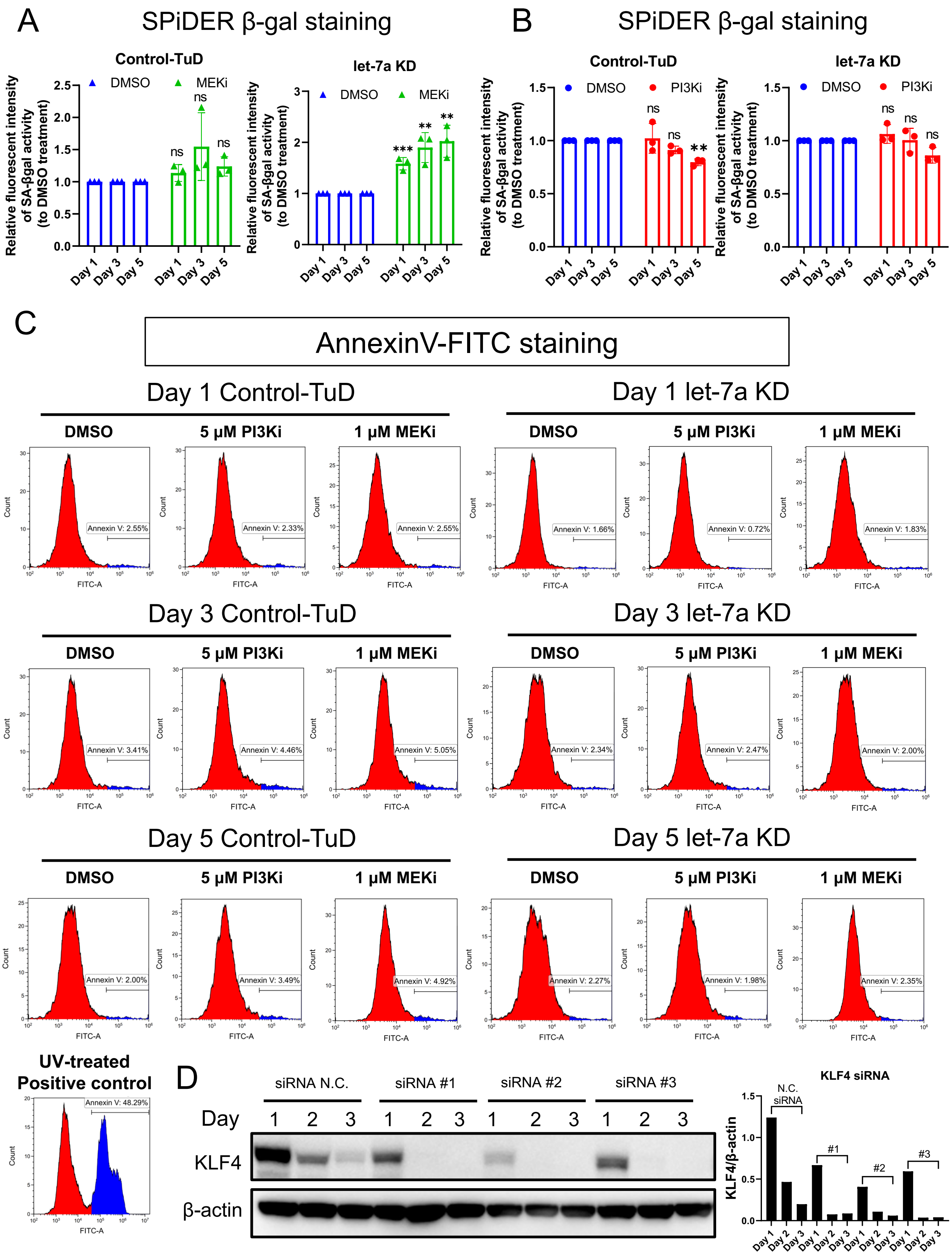

**Supplemental Figure S3. Analysis of senescence and apoptosis.**

(A-B) Bar graph showing the analysis of flow cytometric detection of SA-βgal-expressed cells in control-TuD and let-7a-KD Muse cells (All n=3).

(C) Annexin-V-FITC staining for detecting cellular apoptosis after PI3Ki and MEKi treatment.

(D) Western blot showing the knockdown effect of KLF4 siRNA. β-Actin was used as an endogenous control. The KLF4 expression intensity was normalized by β-actin.

PI3Ki: LY294002. MEKi: PD0325901.

Li_Supplemental_Fig_S4

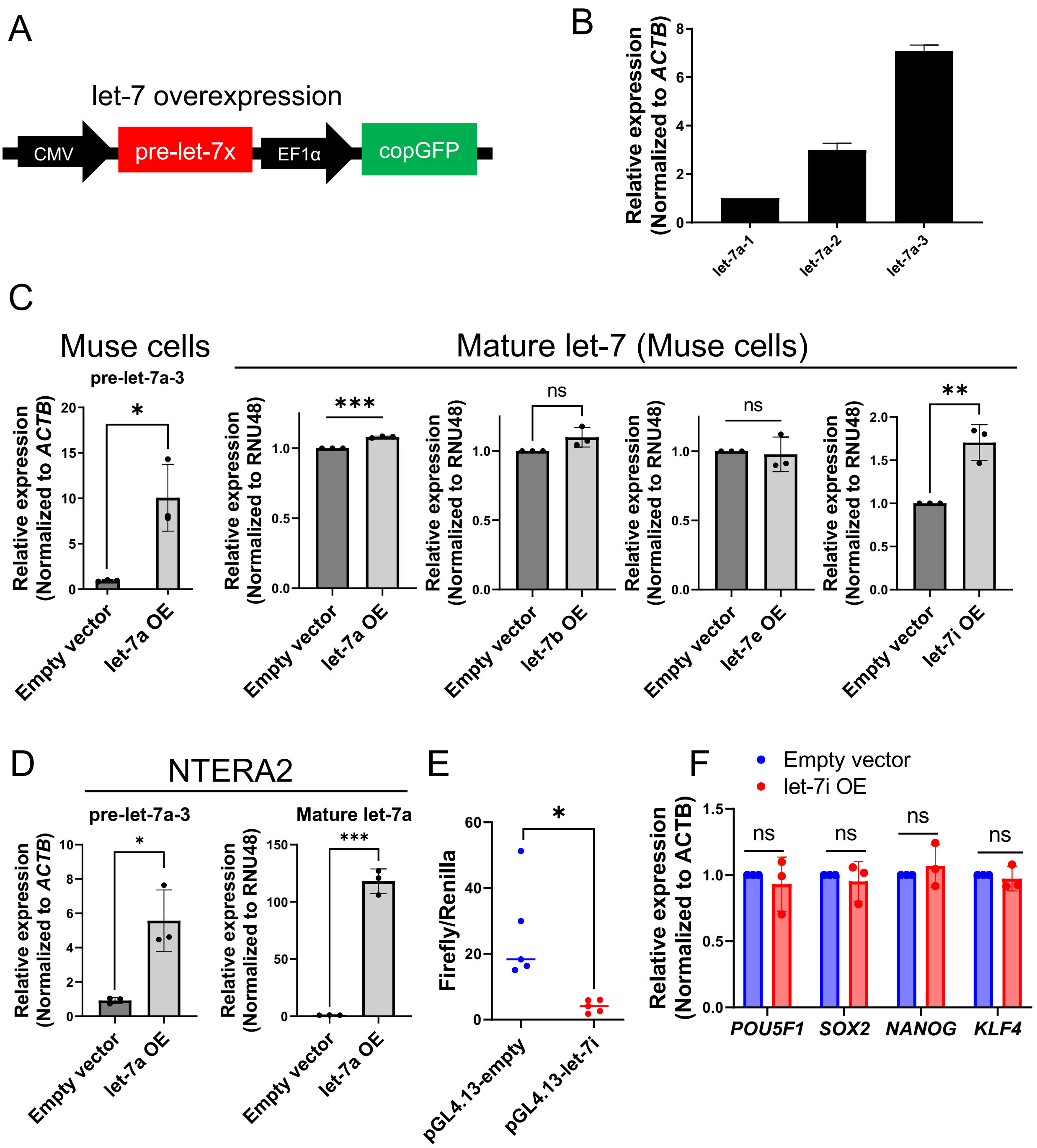

**Supplemental Figure S4. Let-7 overexpression in Muse cells.**

(A) Design of the let-7 overexpression system. copGFP was used as an indicator of positively transfected cells.

(B) Comparison of the expression of let-7a-1, let-7a-2, and let-7a-3 in Muse cells (n=3).

(C) qPCR analysis showing the overexpression of pre-let-7a-3 and mature let-7 (n=3).

(D) qPCR showing the overexpression of pre-let-7a-3 and mature let-7a in NTERA2 (both n=3). Empty vector-transfected cells were used as a negative control.

(E) Luciferase assay to confirm the effect of let-7i overexpression (n=5). Renilla luciferase was used as a control.

(F) qPCR analysis of the expression of pluripotency genes before and after let-7i overexpression (n=3).

OE: overexpression. ACTB and RNU48 were used as endogenous controls for gene expression and miRNA expression, respectively.

Li_Supplemental_Fig_S5

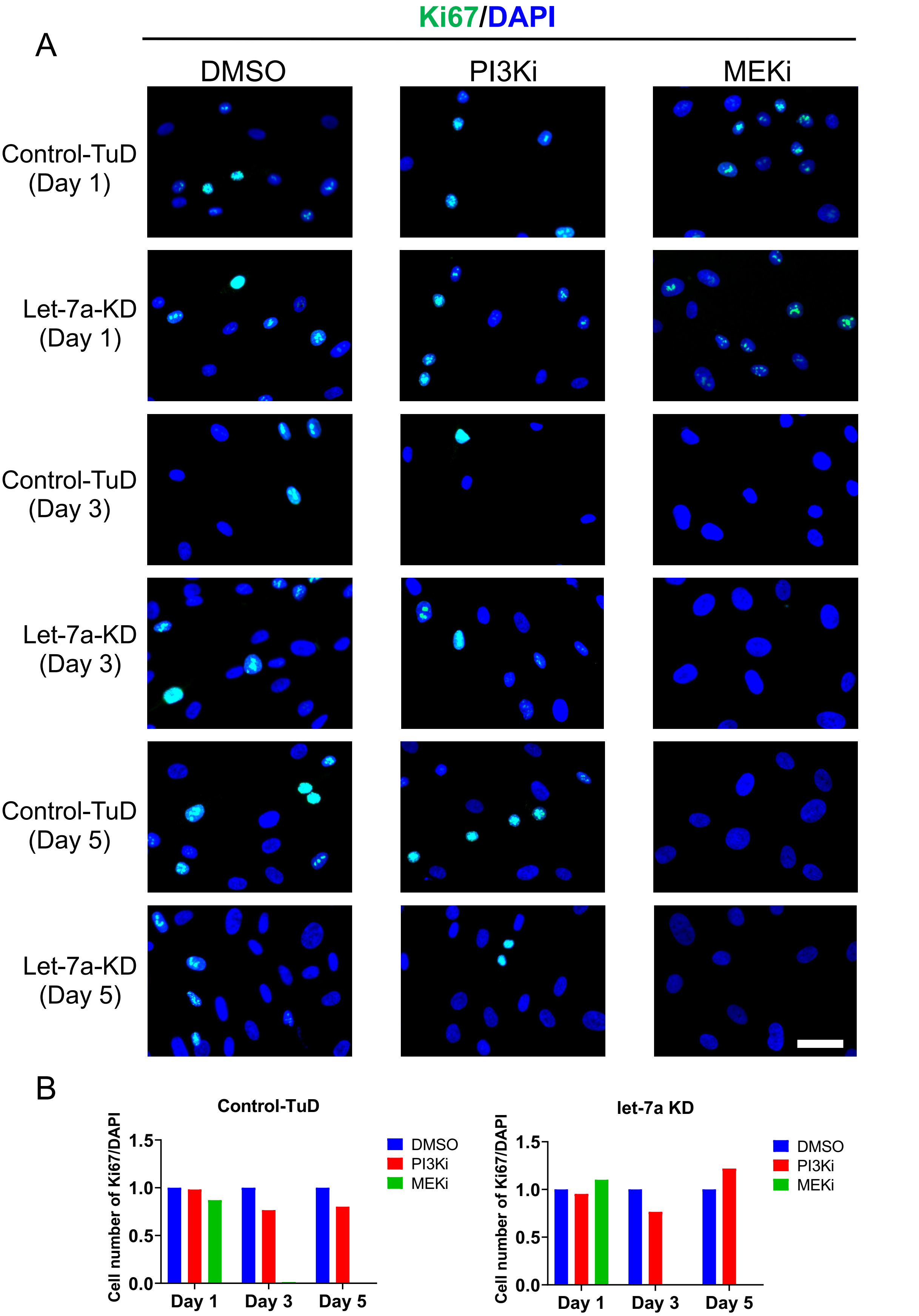

**Supplemental Figure S5. Ki67 staining in DMSO, control-TuD-, and let-7a-KD-Muse cells**

(A) Immunostaining for Ki67. Scale bar: 50 μm.

(B) Cell count ratio of Ki67 to DAPI.

PI3Ki: LY294002. MEKi: PD0325901. Blue: DAPI. Green: Ki67.

**Supplemental Table 1: Primers**

| Name | Sequence (5'→3') or assay name | |
|---|---|---|
| *LIN28A* | Taqman Gene Expression Assays | Hs00702808_s1 |
| *LIN28B* | Taqman Gene Expression Assays | Hs01013729_m1 |
| hsa-let-7a | Taqman MicroRNA Assays | Assay ID: 000377 |
| hsa-let-7b | Taqman MicroRNA Assays | Assay ID: 002619 |
| hsa-let-7c | Taqman MicroRNA Assays | Assay ID: 000379 |
| hsa-let-7d | Taqman MicroRNA Assays | Assay ID: 002283 |
| hsa-let-7e | Taqman MicroRNA Assays | Assay ID: 002406 |
| hsa-let-7f | Taqman MicroRNA Assays | Assay ID: 000382 |
| hsa-let-7g | Taqman MicroRNA Assays | Assay ID: 002282 |
| hsa-let-7i | Taqman MicroRNA Assays | Assay ID: 002221 |
| hsa-mir-98 | Taqman MicroRNA Assays | Assay ID: 000577 |
| RNU48 | Taqman MicroRNA Assays | Assay ID: 001006 |
| hsa-let-7a-1 | Taqman Pri-miRNA Assay | Hs03302533_pri |
| hsa-let-7a-2 | Taqman Pri-miRNA Assay | Hs03302539_pri |
| hsa-let-7a-3 | Taqman Pri-miRNA Assay | Hs03302546_pri |
| *ACTB* | Taqman Gene Expression Assays | Hs03023880_g1 |
| *ACTB*-F | CATGTACGTTGCTATCCAGGC | PrimerBank ID: 4501885a1 |
| *ACTB*-R | CTCCTTAATGTCACGCACGAT | PrimerBank ID: 4501885a1 |
| *POU5F1*-F | AACCCACACTGCAGCAGATCA | NCBI Primer-BLAST |
| *POU5F1*-R | ACACTCGGACCACATCCTTC | NCBI Primer-BLAST |
| *SOX2*-F | TCCAACATCCTGAACCTCAGC | NCBI Primer-BLAST |
| *SOX2*-R | TCTGCGTCACACCATTGCT | NCBI Primer-BLAST |
| *NANOG*-F | CAGCTCGCAGACCTACATGA | NCBI Primer-BLAST |
| *NANOG*-R | CTCGGACTTGACCACCGAAC | NCBI Primer-BLAST |
| *KLF4*-F | CACCCACACTTGTGATTACGC | NCBI Primer-BLAST |
| *KLF4*-R | TGTTTACGGTAGTGCCTGGTC | NCBI Primer-BLAST |

**Supplemental Table 2: Primary antibodies**

| Antibodies | Manufacturer | Catalog number |
|---|---|---|
| Purified anti-human/mouse SSEA-3 Antibody | BioLegend | 330302 |
| Purified Rat IgM, κ Isotype Ctrl Antibody | BioLegend | 400801 |
| Fluorescein (FITC) AffiniPure Goat Anti-Rat IgM, μ chain specific | Jackson ImmunoResearch | 112-095-075 |
| Allophycocyanin (APC) AffiniPure $F(ab')_2$ Fragment Goat Anti-Rat IgM, μ chain specific | Jackson ImmunoResearch | 112-136-075 |
| LIN28A | Cell Signaling | 3978S |
| LIN28B | Cell Signaling | 4196S |
| IGF-I Receptor β | Cell Signaling | 9750 |
| Insulin Receptor β | Cell Signaling | 23413 |
| IRS2 | Abcam | Ab134101 |
| NRAS | Santa Cruz Biotechnology | SC-31 |
| β-actin | Abcam | ab6276 |
| Phospho-AKT (T308) | Cell Signaling | 13038S |
| Phospho-AKT (S473) | Cell Signaling | 9271 |
| AKT | Cell Signaling | 9272 |
| Phospho-p44/p42 MEK/ERK (ERK1/2) (Thr202/Tyr 204) | Cell Signaling | 9101 |
| p44/p42 MEK/ERK (ERK1/2) | Cell Signaling | 9102 |
| KLF4 | Cell Signaling | 12173S |
| Peroxidase AffiniPure Goat Anti-Mouse IgG | Jackson ImmunoResearch | 115-035-071 |
| Peroxidase AffiniPure Goat Anti-Rabbit IgG (H+L) | Jackson ImmunoResearch | 111-035-144 |
| Ki67 | Abcam | Ab16667 |
| Alexa Fluor 488 AffiniPure $F(ab')_2$ Fragment Donkey Anti-Rabbit IgG (H+L) | Jackson ImmunoResearch | 711-546-152 |
